# Insights into the patterns of molecular evolution and functional diversification of *NAC* gene family in land plants

**DOI:** 10.1101/2025.09.22.677750

**Authors:** Yuxin Liu, Zelong Li, Yilu Zhou, Yu Wen, Liyuan Guo, Xiaohong Chen, Zhaoxue Han

**Author notes:** These authors contributed equally to this work (Yuxin Liu, Zelong Li, Yilu Zhou). Corresponding author: Zhaoxue Han.

## Abstract

NAC proteins are involved in various aspects of plant development and stress responses. However, the evolutionary history and diversification patterns of *NAC* genes have not been examined systematically in land plants. By identifying 5052 *NAC* genes in 46 species of land plants and constructing their phylogenies, we found that *NAC* genes fell into three categories of evolutionary clades. An integrative analysis of evolutionary clades and synteny network revealed that *NAC* syntelogs exist widely across angiosperms, and ancient tandem duplications and multiple lineage-specific transposition events contributed to expansion and divergence of *NAC* family. Based on the reconstructed phylogenies, a potential evolutionary framework of *NAC* genes in land plants was presented, indicating that 13 ancestral genes which showed distinct expansion patterns evolved all current *NAC* repertoires of Arabidopsis and maize. *ZmNAC* genes exhibit divergent expression diversification patterns, which are generally in agreement with the expected clade- and ancestral gene specific expression characterization. The subcellular localization analysis of 14 ZmNAC proteins demonstrated that they are predominantly localized in the nucleus. Transcriptional activity analysis revealed their functional divergence in transcriptional activity. Through *ZmNACs* overexpression Arabidopsis, we identified the roles of *ZmNACs* under drought and salt stresses, thereby substantiating functional differentiation of *NAC* transcription factors in abiotic stress responses. Overall, the synteny network analysis and the reconstructed evolutionary framework of *NAC* genes in this study increased our understanding for the evolutionary history and functional consequences of *NAC* genes, and in return, *ZmNAC* expression profiles are also crucial for understanding the evolutionary sequence divergence from a functional perspective. Our results deepen the knowledge of the evolutionary mechanisms of *NAC* genes and the function of maize *NAC* genes in abiotic stress responses.

**Highlights:** Identified 5,052 *NAC* genes across 46 land plant species, revealing a maize comprehensive evolutionary framework with 13 ancestral genes.

Ancient tandem duplications and lineage-specific transposition events drove *NAC* family expansion, as uncovered by synteny network analysis.

Maize *NAC* genes exhibit clade-specific expression divergence, nuclear localization, and functional variability in transcriptional regulation under abiotic stress.

Transgenic Arabidopsis overexpressing *ZmNACs* confirmed their functions in drought and salt stress tolerance, showing functional differentiation among *NAC* clades.

## Introduction

Transcription factors (TFs) act as a class of functionally important regulatory proteins in a variety of biological processes. In plants, NAC (NAM/ATAF/CUC2) proteins constitute one of the largest families of TFs and they are present in a wide range of plants. Since the first genome-wide structure and function study on *NAC* genes in *Arabidopsis thaliana* and *Oryza sativa* (Ooka et al., 2003), *NAC* genes have been surveyed across a large number of plant genomes, most of which belong to land plants. Typically, NAC TFs contain a highly conserved N-terminal DNA-binding NAC domain, which can be divided into five subdomains (A–E), and a variable C-terminal domain (transcriptional regulation region) (Nakashima et al., 2012). Some NAC proteins are designated as NAC membrane-bound TFs (NTL1-like, NTLs) with transmembrane motifs (TMs) (Kim et al., 2006; Seo et al., 2008). NAC TFs primarily bind to the NACRS and play important roles in many biological processes by activating or inhibiting the expression of target genes (He et al., 2018; Yuan et al., 2020).

In the past years, numerous *NAC* genes have been functionally characterized in the model system *Arabidopsis thaliana* and some important crops. The early examples are Arabidopsis *CUC1* and *CUC2* which are involved in organ separation (Hibara et al., 2003; Takada et al., 2001), *ANAC019*, *ANAC055* and *ANAC072* showed a significant increase in drought tolerance (Tran et al., 2004), and *VND-INTERACTING2 (VNI2)* involved in regulating salt tolerance via COR and RD genes (Yang et al., 2011). Some other Arabidopsis *NAC* genes have been implicated in the control of sieve element morphogenesis (Furuta et al., 2014), leaf senescence (Kim et al., 2016), mitochondrial retrograde signaling (Meng et al., 2019; Ng et al., 2013), *ANAC070* transcription factor confers aluminum tolerance in Arabidopsis by repressing the ANAC017-XTH31 module to reduce root cell wall Al accumulation and hemicellulose/xyloglucan content, while potentially modulating the STOP1-mediated Al tolerance pathway(Li et al., 2024), The root cap-localized *NAC* transcription factor *SOMBRERO (SMB)* mediates root halotropism by establishing spatiotemporal auxin asymmetry in the lateral root cap (LRC) through the basal regulation of the auxin influx carrier *AUX1*, enabling directional root bending away from saline environments(Zheng et al., 2024). In crops, previous research has indicated that *SNAC1* and *SNAC2* enhance abiotic stress tolerance in transgenic rice (Hu et al., 2006, 2008), *OsNAC3* positively regulates rice seed germination by directly activating the ABA catabolic gene *OsABA8ox1*(Huang et al., 2024). *OsNACIP6* confers drought tolerance in rice by binding to the *ABRE4* cis-element to upregulate *OsNAC78* expression, which activates downstream glutathione reductase gene *OsGSTU37*, forming the *OsNACIP6/OsNAC78-OsGSTU37* transcriptional module that enhances oxidative stress resistance (Yu, et al., 2024). Several rice *NAC* genes have been shown to influence biotic stress response (Sun et al., 2013; Yoshii et al., 2010), abiotic stress tolerance (Fang et al., 2015; X. Zheng et al., 2009), and root development (Mao et al., 2019). Wheat *TaNAC29* improved salt stress tolerance through enhancing the antioxidant system and regulating the abiotic stress-response(Xu et al., 2015), Ta*NAC2* interacts antagonistically with TaLBD41 to regulate wheat adaptation to soil nitrate by competitively binding to key nitrogen uptake/reduction genes (*TaNRT2.1, TaNR1.2, TaNADH-GOGAT*)(Jing et al., 2024). In maize, a prominent example is *ZmNAC111*, which enhanced tolerance to dehydration (Mao et al., 2015); *ZmNAC20* enhances maize drought resistance by promoting ABA-mediated stomatal closure to reduce water loss and activating stress-responsive gene expression, resulting in elevated leaf water retention and plant survival under drought conditions(Liu et al., 2023). *ZmNAC128* and *ZmNAC130* cooperatively regulate maize endosperm filling by directly activating genes involved in nutrient transport (including *ZmSWEET4c*, *ZmSUGCAR1* and *ZmYSL2*) and zein biosynthesis (γ-zeins)(Chen et al., 2023), several other several maize *NAC* genes are mainly involved in antioxidant defense (Zhu et al., 2016), endosperm development, and grain yields (Zhang et al., 2019), etc. Recent studies. These studies have linked NAC proteins to a variety of aspects of biological processes, including plant growth, development and stress responses. Despite the biological importance of the *NAC* family, the functions of most members within this family remain unknown in both model and non-model plants. Large-scale and detailed evolutionary analysis can provide robust insights for functional analysis. While the evolution of *NAC* genes has been frequently investigated based on genome sequences from a single or a limited number of species (Baranwal & Khurana, 2016; Hussey et al., 2015; Jin et al., 2017; Kim et al., 2025; Mohanta et al., 2020; Zhu et al., 2012), there is still a limitation in understanding the evolutionary fate and molecular functions of *NAC* gene family among species from multiple lineages. Benefiting from the recent availability of more and more high-quality plant genome sequences, more comprehensive studies have become feasible to gain new insights into the possible evolutionary mechanism of *NAC* TFs across multiple species and thus shed lights on the distinct functional features of the family.

In this study, we expanded the species scope of previous research by identifying members of the *NAC* family across 46 species. Subsequently, we examined tandem duplication events among land plant *NAC* family members, revealing the contribution of tandem duplication to the expansion of different *NAC* clades. By constructing synteny networks across these 46 species, we further validated the topological structure of phylogenetic trees and elucidated the expansion patterns and evolutionary history of the *NAC* family in land plants. The *NAC* family underwent large-scale expansion after the emergence of angiosperms and experienced lineage-specific expansions around speciation events. Furthermore, we constructed evolutionary frameworks for *NAC* genes acrossdifferent lineages of land plants, revealing divergent evolutionary trajectories and distinct fates of ancestral *NAC* genes in different species.

Using maize *NACs* as an example, we demonstrated the differential fates of *ZmNACs* derived from distinct ancestral genes at the transcriptional level through tissue-specific expression profiling and drought stress response analysis. Subcellular localization and transcriptional activity assays at the protein level confirmed both functional conservation and divergence among *ZmNAC* transcription factors of different ancestral origins. By generating Arabidopsis overexpression lines for *ZmNACs*, we identified five *ZmNACs* with functional roles in drought and salt stress responses, further demonstrating the functional diversity of *NAC* transcription factors in regulating plant stress responses. Our study conducted functional investigations of *NAC* transcription factors to determine their specific roles, while elucidating their contributions to plant development and stress responses.

Our work provides a reference for further understanding of the evolutionary trajectories of the *NAC* genes and discovering potential *NAC* functional genes in maize and other crops.

## Results

### The evolutionary history of NAC transcription factors in land plants

To comprehensively identify NAC proteins in green plants, based on the conservation of HMM profiles of NAC proteins, we performed a genome-wide sequence homology search and identified 5,052 putative *NAC* genes, with variable copy number distribution across 46 genomes investigated (Fig. 1; Table S1). Most seed plants have more than 70 *NAC* genes and the average number in seed plants is 113. In angiosperms, the copy number of *NAC* genes varies considerably, ranging from 48 in *Beta vulgaris* to 226 in *Malus x domestica*. In contrast, two early-diverged land plants *Selaginella moellendorffii* and *Physcomitrella patens*, contain only 20 and 30 *NAC* genes respectively, though *Physcomitrella patens* experienced an ancient PHPA-α whole genome duplication (WGD) (Leebens-Mack et al., 2019) (Fig. 1; Table S1). Therefore, we inferred that the extensive expansion of *NAC* genes happened after the divergence of seed plants, accompanying with numerous whole genome polyploidy events. This inference is also consistent with the result of Alejandro Pereira-Santana *et al*. (Leebens-Mack et al., 2019); Leebens-Mack *et al*. (Leebens-Mack et al., 2019)

**Figure. 1.**
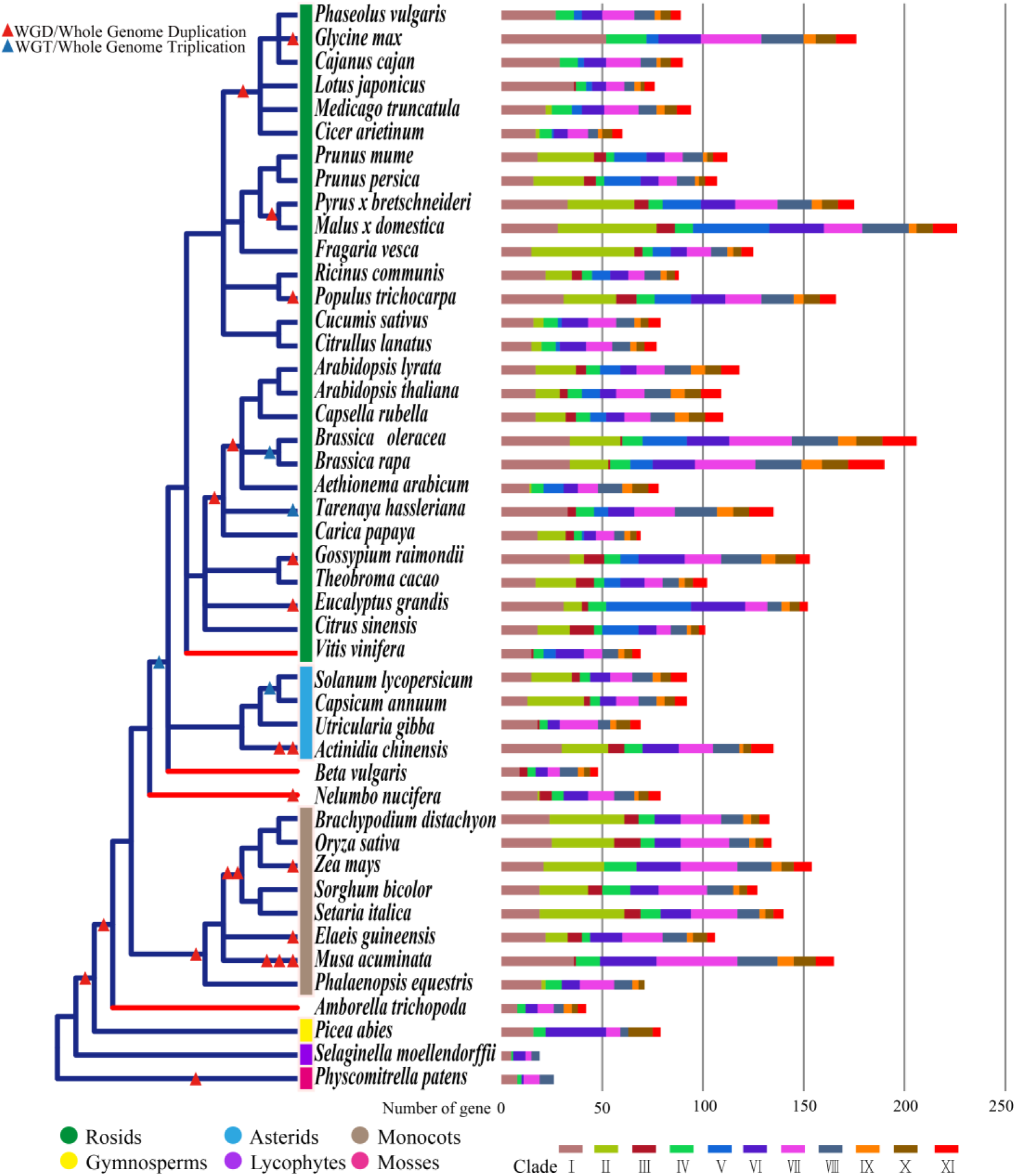
Species phylogeny and number of *NAC* family members in 46 species. The phylogenetic tree of 46 species was inferred using NCBI Taxonomy Browser. The red branches in the phylogenetic tree indicate the basal rosid *Vitis vinifera*, the basal eudicots *Beta vulgaris* and *Nelumbo nucifera*, and the basal angiosperm *Amborella trichopoda*, respectively. Different color bars which are on the left of species names represent different species categories. The right bars of species names indicate the number of *NAC* genes in each species and the distribution in each clade with a specific color bar at the bottom (see Supporting Information Table S1 and Fig. S1). The red and blue triangle on the phylogenetic tree indicate whole-genome duplication (WGD), and whole-genome triplication (WGT), respectively.

To explore the evolution of the *NAC* gene family in plants, a phylogenetic tree was constructed using full-length sequences of all 5,052 NAC proteins from 46 plant species. as shown in Fig. S1 and Table S2, all *NAC* genes except eight ones were clustered into 11 clades (named as Ⅰ, Ⅱ, …, Ⅺ). Each clade all contains members from angiosperms (Fig. S1; Tables S1, S2). Clades Ⅰ, Ⅳ, Ⅵ, Ⅶ and Ⅷ are land plant-specific, consisting of members from mosses, lycophytes, gymnosperms, and angiosperms, so these clades may originate from mosses or earlier species. Clades Ⅹ and Ⅺ just contain members from gymnosperms and angiosperms (Fig. S1; Tables S2, S3), indicating that these two seed plant-specific clades are derived from ancestral genes that predate the divergence of gymnosperms. Clades Ⅱ, Ⅲ, Ⅴ and Ⅸ are angiosperm-specific (Fig. S1), suggesting that these four clades occurred recently and may form from angiosperms. Interestingly, the *NACs* in Clade Ⅴ are rosid-specific (Table S2), suggesting that the genes in Clade Ⅴ may be the youngest ones in this study.

Furthermore, the number of *NAC* genes across different lineages within different clades is also divergent. For instance, the number of *NAC* genes of the legume family is relatively less among the four angiosperm-specific clades, with the highest count of 12 members (12/94, 12.77%) exist in *Medicago truncatula* of the legume family. By contrast, within the Rosaceae family, *Fragaria vesca* has the highest number of 67 members (67/125, 53.6%) (Table S2). These results indicate that the *NAC* gene family likely underwent lineage-specifc expansion following the differentiation of seed plants. Furthermore, the distinctive signatures in the conserved motifs of the NAC domain significantly differentiate the 11 clades. Through sequence analysis of the five subdomains within NAC domains across different clades, we found that the NAC domains in land plant-specific and seed plant-specific clades are more conserved compared to those in angiosperm-specific clades. Particularly, clades II, III, and V demonstrate higher motif diversity in subdomains C and D(Fig. S2).

### Divergent conservation of tandem duplications associated with differential *NAC* clades expansion in land plants

Gene duplication is an important mechanism for acquiring new genes and creating genetic novelty in organisms. The duplications of genomic content occur on various scales by independent mechanisms, including tandem duplications, segmental duplications and whole-genome duplications (Ramsey & Schemske, 1998; Wendel & Flagel, 2009). We investigated the physical positions of 5,052 *NAC* genes on each chromosome across 46 species, and finally 1,178 tandem duplications, 716 tandem pairs, and 463 tandem groups were identified (Table S5). The number of tandem duplications is unevenly distributed in different clades of the phylogenetic tree. Within Clades Ⅱ, Ⅴ and Ⅺ, there are 38.18% (247/647), 76.11% (239/314) and 38.31% (113/295) tandem duplications, respectively. This suggests that tandem duplication events greatly contribute to the expansion of these clades, especially in the rosid-specific Clade Ⅴ. This is similar with the DUF295 family, which underwent numerous rounds of tandem duplications in *Brassicaceae* (Lama et al., 2019), indicating that lineage-specific tandem duplications are the main contributor of gene family rapid expansion in a short period of time. And more, the number of tandem duplications in different species is also diverse. In monocots, the ratio ranges from 12.68% (9/71) in *P. equestris* to 26.67% (36/135) in *O. sativa*, while in eudicots, the proportion of tandem duplications ranges from 6.52% (6/92) in *C. annuum* to 53.95% (73/152) in *E. grandis* genome (Table S5).

The partners of most tandem duplications have high sequence similarity and they are distributed closely in the phylogenetic tree. While some tandem partners are far apart in the phylogenetic tree, even are clustered in different clades (Fig. 2a). According to the distribution of tandem duplications across phylogenetic clades, we further categorized the tandems into two types, including Clade-Specific (CS) tandems (each pair is located in same clade), and Clade-non-Specific (CnS) ones (the partner of each tandem pair are located in different clades, respectively). Within the identified 716 pairs of tandem duplications, 537 pairs were CS type and 179 pairs were CnS type (Table S5). We observed that 93.72% (224/239) in Clade V and 91.5% (226/247) in Clade Ⅱ of tandem duplications are CS type, while only 13.46% (14/104) and 29.17% (7/24) of tandem duplications are CS type in Clade Ⅶ and Clade Ⅸ, respectively (Table S5). We then noticed that there are 66 pairs of CnS type tandems being widespread in angiosperms between Clade Ⅰ and Clade Ⅶ, with most of the partners are clustered together in each clade. Similarly, there are 15 pairs of CnS-tandems between Clade Ⅵ and Clade Ⅸ, 13 pairs of CnS-tandems between Clade Ⅰ and Clade Ⅺ (Fig. 2a; Table S5), etc. The divergence in the number and types of tandem duplications across different species and clades indicates the evolutionary complexity of tandem duplication events. Genes from ancient tandem duplications are not conserved in most evolutionary lineages, while tandem duplication genes from more recent origins are conserved within evolutionary branches.

**Figure. 2.**
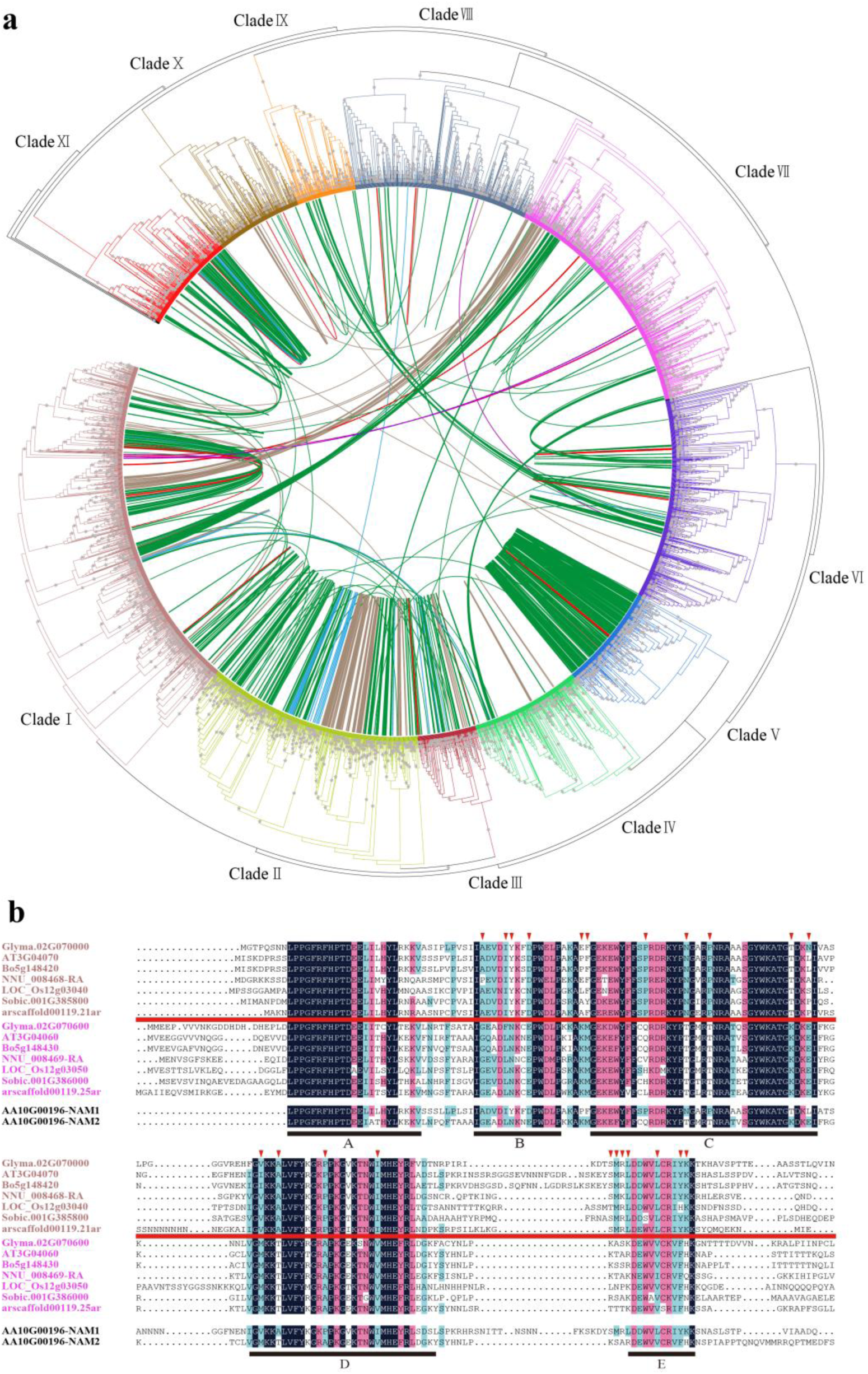
Tandem duplications of NAC genes in 46 plants. (a) Phylogenetic location of tandem duplications in 46 species. Tandem duplications are connected by lines. The colors of lines are marked according to the species phylogeny (Table S5 and Fig. 1), gray stars on the branches indicate bootstrap support values (>0.85). (b) The polypeptide sequence align of the representative *NAC* tandem duplications between Clade Ⅰ and Clade Ⅶ. The five conserved *NAC* subdomains are underlined. The conserved divergent amino-acid sites are marked by inverted triangles.

To validate whether the evolutionary time differences caused the CS and CnS tandem types or not, we further analyzed NAC domain sequences of the CS in Clade V and CnS-tandem partners between Clade Ⅰ and Clade Ⅶ (Fig. 2a; Table S5). Interestingly, most of the CnS-tandem partners in each clade exhibited significantly similar divergent patterns at some amino-acid sites, (Fig. 2b). Therefore, we speculate that the differentiation of these CnS-tandems between different clades is likely to be caused by tandem duplication events occurring in an ancestral organism. The sequences of these tandems within the ancestral species produced variation and then were retained during the evolution process.

### Angiosperm-specific syntenic relationships of *NAC* genes

A broad synteny network analysis is a powerful approach to understand the evolutionary trajectory of genes and genomes both inter-species and intra-species, especially for gene families (Zhao et al., 2017; Zhao & Schranz, 2017). To explore the evolutionary history and the expansion or loss events of *NAC* genes across different lineages of land plants, we employed a synteny network approach to analyze all protein models from 46 genomes for all possible intra- and interspecies whole genome comparisons.

We built a database that contains all the links between syntenic gene pairs existing in syntenic genomic blocks. This database contains in total 917,633 nodes (i.e., genes that were connected by synteny with another gene) and 9,362,666 edges (i.e., pairwise syntenic connections). We then extracted the synteny sub-network of the 5,052 identified *NAC* genes from the entire network database. This sub-network contained 3,783 nodes that were linked by 48,513 syntenic edges (Fig. 3a; Table S6). We found that *NAC* syntenic relationships are widely present across angiosperm lineages, and there are no *NAC* syntelogs in *P. abies* (Gymnosperm). In *S. moellendorffii* (Lycophytes) and *P. patens* (Moss), *NAC* syntelogs were only present in intra-species (Table S6). This is likely due to the fragmented early-version genome assemblies and the extreme phylogenetic distance or less species selected in this study. We visualized this sub-network using Gephi, with nodes color-coded according to the phylogenetic tree (Fig. 4a). We then used the CytoCluster to cluster the synteny sub-networks and obtained 95 communities (Fig. 3a, Table S7).

**Figure. 3.**
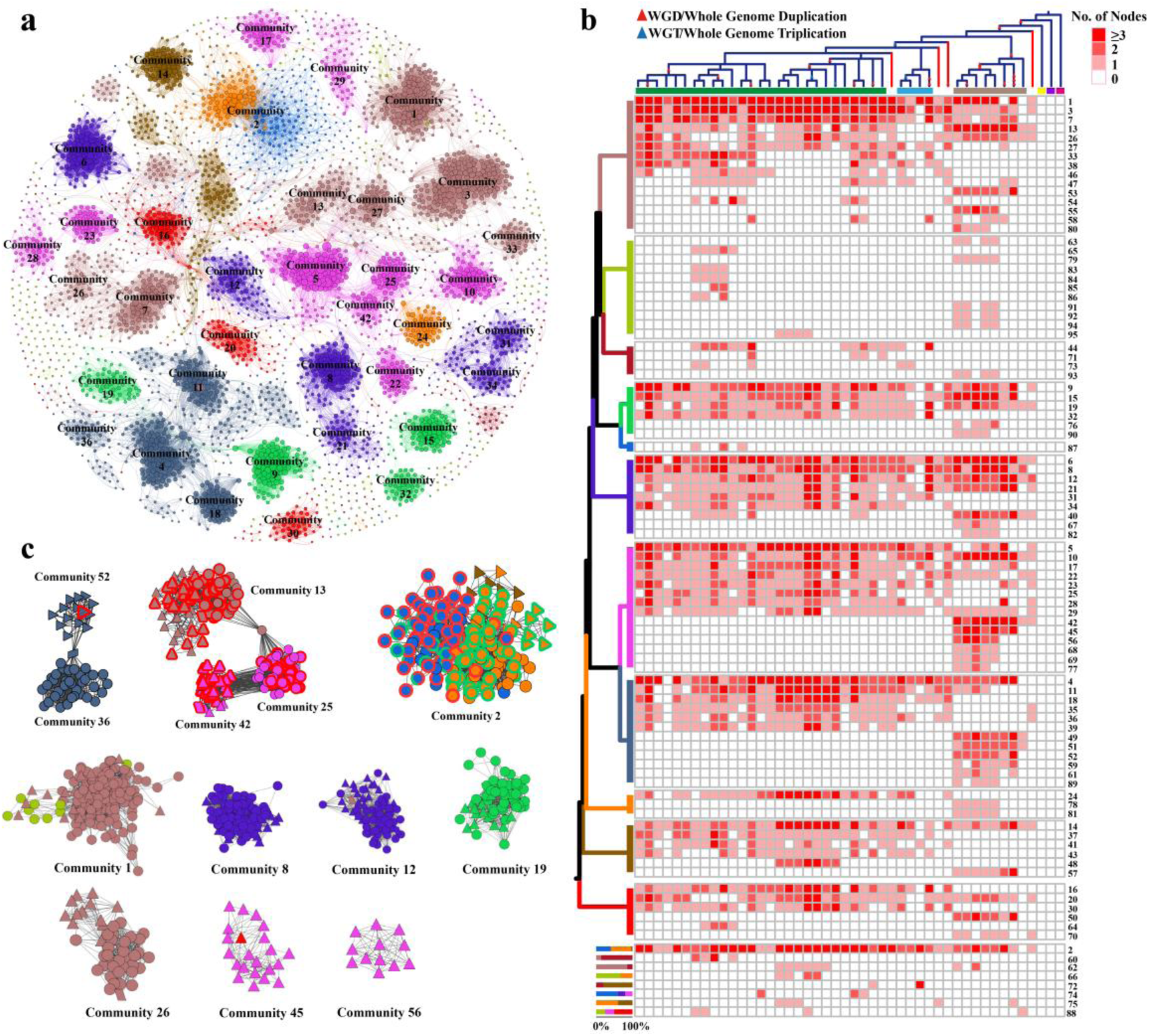
Phylogenomic synteny network of the *NAC* gene family among angiosperms. (a) The syntenic relations of *NAC* genes in the synteny network. The size of each node corresponds to the number of edges it has (node degree). The color of nodes represents different clades and corresponds to Fig. 2. Communities were labeled based on the size number of nodes involved. (b) Phylogenetic profile showing the number and distribution of the syntenic *NAC* genes in plants. Rows represent synteny communities and columns indicate species. The species tree on top of the profile was ordered from the most recent to the most ancient, from the left to the right and corresponds to Fig. 1. The colors on left of the profile represent different clades and correspond to communities are at the bottom. (c) Close-up of the networks for representative communities from Fig. 3a. Node shapes represent different species categories, the circle represents dicots, the triangle represents monocots, and the parallelogram represents the basic angiosperm. Node colors represent different clades. The red color of node border represents tandemly duplicated genes and green color represents *NTL* genes.

**Figure. 4.**
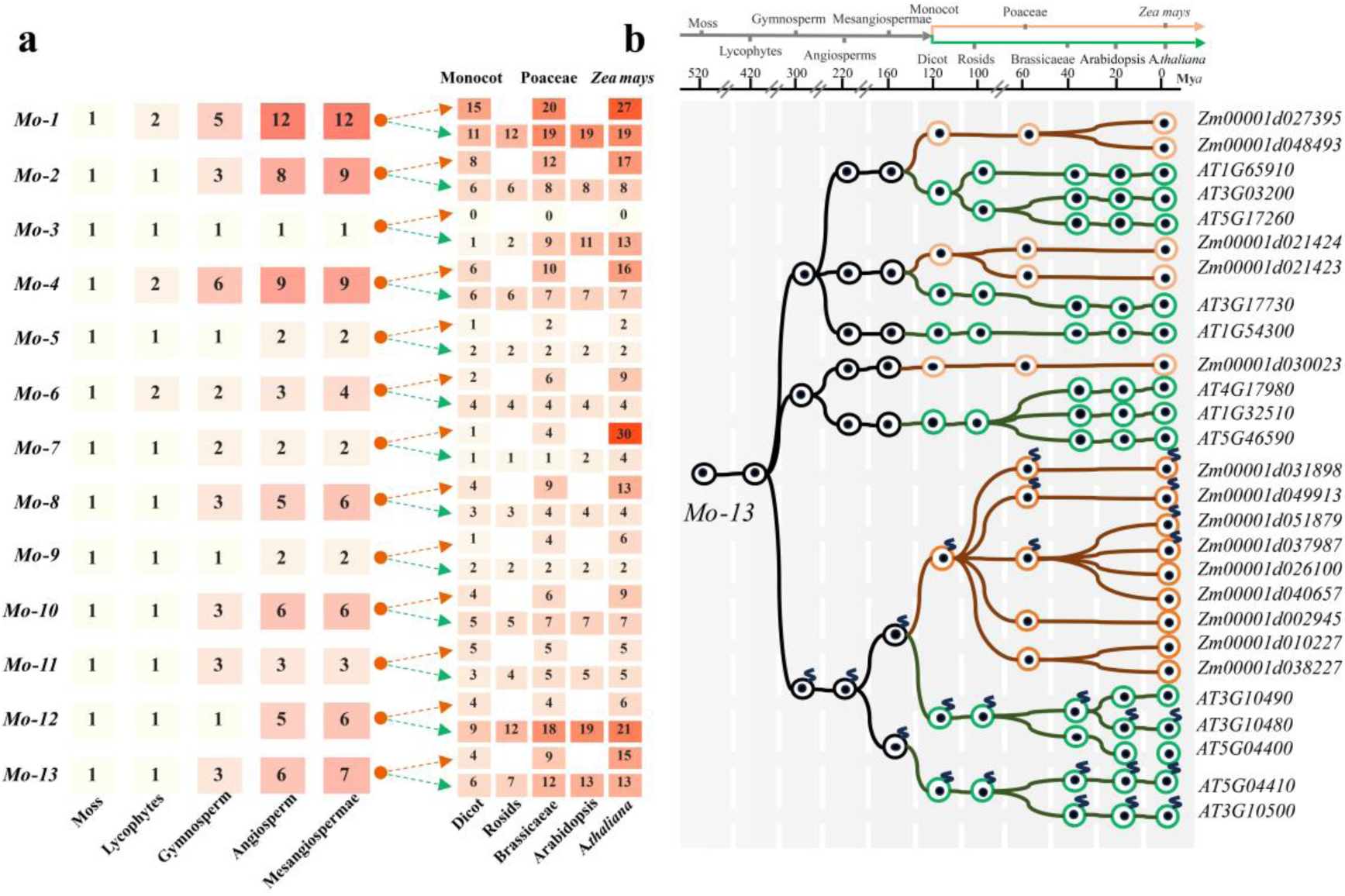
The framework of Arabidopsis and Maize ancestral *NAC* genes in different lineages and expansion of *NAC* genes from *Mo-13*. (a) The nodes along the timeline represent species divergence times. Evolutionary states of Arabidopsis and maize at distinct phylogenetic nodes were retrieved from the reconstructed *NAC* gene phylogeny, and the heatmap illustrates the number of extant ancestral genes across different evolutionary nodes. (b) The ancestral gene *Mo-13* that is responsible for generating the current Arabidopsis and Maize *NAC* repertoire via gradual expansion. The ancient states of Arabidopsis and Maize at different evolutionary nodes were retrieved from the constructed *NAC* phylogenies. Black helix indicates membrane-bound domain.

In the 95 communities, 61 are lineage-specific communities, including 29 monocot and 32 eudicot-specific communities. This indicates that the syntenic relationships within these lineage-specific communities may occur after the divergence of monocots and eudicots. The rest 34 communities are widespread across angiosperms (at least containing one monocot and one eudicot, or one of lineages is *A. trichopoda*) (Fig. 3b; Table S7).

We noticed that the monocot-specific community 52 in Clade Ⅷ is comprised of homologs of seven monocots except *P. equestris* (Fig. 3c). Interestingly, most syntelogs of community 52 link the one (*arscaffold00017.45ar*) of basal angiosperm *A. trichopoda* in community 36 containing homologs from *A. trichopoda* and eudicots (Fig. 3c; Table S6, S7). This synteny pattern suggests that a gene transposition, or a genomic rearrangement event, or genome fractionation process may have acted on the ancestral *NAC* gene after the split of monocots and eudicots. Furthermore, the ancestral *NAC* gene may lose in *P. equestris* and some eudicots (Fig. 3b). Similarly, community 45 is also monocot-specific, and it also consists of homologs from seven monocots, but this distinct synteny community does not have syntenic relationships with other communities. This may be attributed to a specific transposition event and/or the ancient τWGD shared by all monocots (Jiao et al., 2014). Additionally, the community 85 in Clade Ⅱ is eudicot-specific and contains four syntelogs from apple (*Malus x domestica*) and pear (*Pyrus x bretschneideri*) (Fig. 3b), which is likely due to a specific transposition event and/or the WGD happening in apple and pear (Velasco et al., 2010).

Except lineage-specific syntenic relationships, we also explored ancient tandem duplication events from the synteny network. Of all 1,178 tandem duplications, 60.78% (716/1,178) tandems are in the synteny network. This result suggests that the tandems may happen in early ancestral species or come from polyploidy events (Table S5; Fig. S3). Although the number of CS-tandems is far more than that of CnS type, the syntenic relationships of CS-tandems are far less than that of CnS type. This indicates the CS-tandems were mainly retained in a species-specific manner, and their tandem duplications may occur after lineage split, while CnS-tandems were more probably inherited from ancestral tandemly duplicated genes. For instance, we noticed that 47 pairs among the 66 CnS-tandems (between CladesⅠand Ⅶ) were present in the synteny network, and they are distributed in communities 13, 25 and 42 (Fig. 4c). The 47 tandem pairs are widespread across angiosperms (Fig. S3), and the CnS-tandems in *A. trichopoda* exihibited extensive synteny with other tandem duplications in these three communities. Thus we speculated that most of these CnS-tandems may originate from *A. trichopoda* or earlier ancestral species and more than one pair of CnS-tandems was observed in many species, especially in monocots, which are likely due to recent WGD events in these species (Fig. S3). Interestingly, community 25 and community 13 are linked by a *NAC* gene (*AA10G00196*) from *A. arabicum*. We noticed that this gene contains two NAC domains, and the differential residues of these two domains are consistent with those of syntenic CnS-tandem partners in these 3 communities (Figs. 3b, 4c). This indicates that ancestral tandemly duplicated gene pairs may have been retained in *A. arabicum*, but later these two genes gradually fused into one. For instance, the syntelogs of community 2 are mainly derived from two angiosperm-specific Clades Ⅴ and Ⅸ and show extensive syntenic relationships (Fig. 4c). Furthermore, we noticed that there are 52 tandem duplications in Clade Ⅴ and 58 *NTLs* in Clade Ⅸ, and also, these tandems and *NTLs* are highly connected. Based on the syntenic and phylogenetic relationships in this community, we proposed an evolutionary model where one *NTL* from Clade Ⅸ occurred sequence divergence after the split of rosids family and later got tandemly duplicated. Then, this tandem arrangement underwent multiple rounds of polyploidy events accompanying transmembrane motifs loss and sequence divergence, and eventually evolved to the modern *NAC* genes of Clade Ⅴ (Fig. S4). This indicates that the syntelogs from rosids-specific Clade Ⅴ may diverge from Clade Ⅸ. Subsequently, syntelogs in these two clades were independently evolved and have conserved intra- and interspecies syntenic relationships.

### Restoring the evolutionary history of *NAC* genes in Arabidopsis and maize

To further reveal the evolution details of *NAC* genes in land plants, we took two representative species, *A. thaliana* and *Z. mays,* as examples to construct an evolutionary framework. Additionally, the conserved and divergent expression patterns of maize *NAC* genes were characterized across different clades. We adopted a step-by-step strategy (Shao et al., 2014) to establish an evolutionary framework for Arabidopsis and maize, which could help us comprehensively explain the evolutionary mechanism of *NAC* genes and gain a deeper understanding of their functionary diversification. Multiple plant lineages, Arabidopsis (*A. lyrata and A. thaliana*), Brassicaceae (*B. oleracea*), Rosid (*V. vinifera*), Solanaceae (*S.lycopersicum*), Poaceae (*O. sativa*, *B. distachyon*, *Z. mays*, *S. bicolor*, and *S. italic*), Monocots (*P. equestris*), Basal angiosperm (*A. trichopoda*), Gymnosperm (*P. abies*), Lycophytes (*S. moellendorffii*), and Moss (*P. patens*), were examined to elucidate the evolution trajectory of the *NAC* genes in Arabidopsis and maize (Figs S5-S12). A total of 13 ancestral *NAC* genes were identified in the ancestral lineage of Moss (Mo). The number of genes at each time node represents the quantity of ancestral genes recovered at that specific time node. (Fig. 4a,Table S8).

The evolutionary patterns of *Mo-NAC* genes showed three distinct types: conserved-, gradual-, and rapid expansion. The ancestral *Mo-5* exhibited high conservation in gene copy numbers during evolution. In contrast, the ancestral lineages *Mo-1*, *Mo-10*, *Mo-11* and *Mo-13* demonstrated gradual expansion during evolutionary processes, a trend that persisted following the divergence of monocots and dicots. Ancestral *Mo-2*, *Mo-4*, *Mo-6, Mo-8* and *Mo-9* displayed progressive expansion in monocots, while maintaining conserved copy numbers in dicots following the monocot-dicot divergence. Conversely, *Mo-12* exhibited numerical conservation in monocots but underwent gradual expansion in dicots. The lineage-specific expansion of ancestral *Mo-7* derivatives occurred predominantly through rapid tandem duplication events following divergence from Poaceae. The ancestral *Mo-3* lineage was inherited exclusively from the Arabidopsis genome, with most *NAC* derivatives clustered within angiosperm-specific Clade II. Notably, one *NAC* gene resides in land plant-specific Clade I, suggesting that *Mo-3* likely originated in early land plants and was subsequently lost in maize during evolution.

By tracing the evolutionary history of ancestral *NAC* genes, we discovered that the common ancestral genes of Arabidopsis and maize underwent distinct evolutionary processes in monocots and dicots, reflecting the evolutionary diversity of the *NAC* family. Based on the evolutionary framework of *NACs*, we can gain a deeper view of the evolutionary process of *NACs* within specific species. For example, we identified 8 and 11 *NTLs* in *A. thaliana* and *Z. mays*, respectively. These *NTLs* were from both ancestral *Mo-12* and *Mo-13*. We further traced the gain and loss of the TM of the *NTLs* inherited from *Mo-13*. There are 4 and 3 *NTLs* originating from *Mo-13* in maize and Arabidopsis, respectively (Fig 4b). The *NTLs* of other species from *Mo-13* were assembled on the conserved branches of different evolutionary nodes (Figs S5-S12). We only found one *NTL* gene in Norway spruce (*P. abies*), and the origin of this gene could be traced to the divergence node of the common ancestor gymnosperm in *Mo-13*. However, 5 *NTLs* from a moss species (*P. patens*) were not inherited from the *Mo-13* (Fig S5; Table S8), which indicates all *NTLs* of the *Mo-13* derivatives inherited the TM motif from the *NTL* of *P. abies*. Thus, we inferred that there may be independent evolutionary pathways of the *NTLs* between *P. abies* and *P. patens* (Fig 5A, S5-S12).

**Figure. 5.**
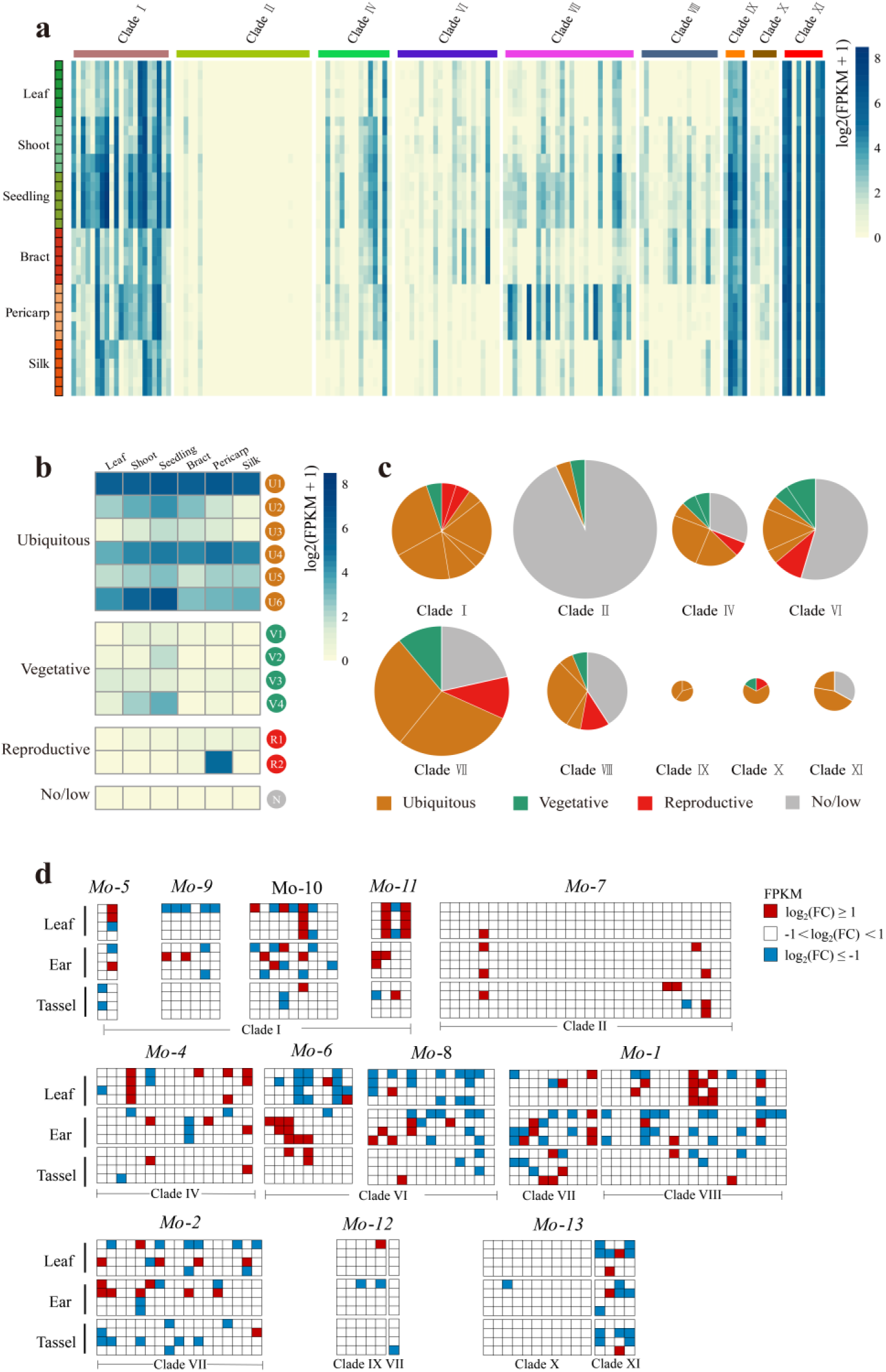
Expression patterns of maize *NAC* genes in diverse tissues and under drought stress. All maize *NAC* genes are shown by clades and expression data (Wang et al. 2018) for all 36 samples are available in Table S10. (a) A heatmap shows expression level of all *ZmNAC* genes in different clades (columns) and maize developmental stages/tissues (rows). Genes and tissues are listed in Tables S10. (b) Mean expression values for the different modules are represented as a heatmap. Colors indicate different general expression patterns: ubiquitous (orange), vegetative (green), reproductive (red) and no or low expression (gray). (c) Different expression modules mapped to the maize *NAC* genes in different clades. The diameter of circles indicated the number of genes in different clades.(d) Response of *ZmNACs* genes to drought stress at different time points.

### Expression behavior of maize *NAC* genes during development and drought stress shows functional diversification

Based on the RNA-seq data of six maize tissues (Wang et al., 2018), We further delineated the expression atlas of all *ZmNACs* in maize six tissues. Ninty-three out of 155 *ZmNACs* were expressed (FPKM>1) in at least one developmental stage, while the other ones showed very low or no expression (No/low) (Fig. 5a; Table S10). We noticed that 86 out of the 94 expressed genes (91.48%) are syntelogs, on the contrary, only 24 out of the 61 No/low genes are syntelogs (39.34%). This result suggests that most syntelogs were preserved and may perform functions in plant developmental stages. Next, the *ZmNACs* were clustered according to their expression patterns. The clustering result was grouped into four expression modules based on expression similarities in vegetative and reproductive tissues (Fig. 5b). The ubiquitous type contained 6 modules (U1-U6), indicating that the genes were expressed in at least one vegetative and one reproductive tissue, respectively. The genes exclusively expressed in vegetative or reproductive tissues were defined as vegetative (V1-V4) or reproductive (R1-R2) types. The remaining genes which showed a very low expression or no expression were considered as No/low type (FPKM < 1) (Fig. 5b; Table S10).

After clustering, it was observed that different clades exhibited distinct expression characteristics, Most *ZmNACs* in Clade I showed constitutive expression; Most genes in Clade II displayed low/no expression. Clades IV, VI, VII, and VIII showed significant differentiation: some were constitutively expressed, some were expressed in vegetative/reproductive tissues, and some showed low/no expression; Clade IX contained a small number of *ZmNACs*, all of which were constitutively expressed; In Clade X, some *ZmNACs* were constitutively expressed, while others were expressed in vegetative and reproductive tissues; Clade XI included *ZmNACs* with either constitutive expression or low/no expression. This result revealed marked divergence in expression patterns between different clades, with varying degrees of differentiation within each clade, reflecting the functional diversity of *ZmNACs* in regulating maize growth and development (Fig. 5c).

Many *NAC* genes are considered to be responsive to abiotic stress. To explore the involvement of *NAC* genes in drought stress responses, we analyzed RNA-seq data from three tissues (leaf, ear, and tassel) of maize B73 at four developmental stages (V12, V14, V18, and R1) under both well-watered and drought stress conditions. Using FPKM values, we constructed expression profiles to investigate the expression patterns of maize *NAC* genes under drought stress (Fig. 5d). 40% (62/155) of *NAC* genes respond to drought stress, and most exhibit collinear relationships. In Clade II, 83% (25/30) of *NAC* genes do not respond to drought stress, which aligns with tissue-specific expression analysis. We classified *ZmNACs* by ancestral genes to investigate their stress responses: *Mo-1,* consisting of Clade VII and VIII, preferentially responding to drought stress in leaf and ear tissues; *Mo-2,* consisting of Clade VII, exhibiting similar drought response levels across three tissues; *Mo-4,* consisting of Clade IV, preferentially responding to stress in leaf and ear tissues; *Mo-5,* consisting of Clade I, with all members responding to stress; *Mo-6,* consisting of Clade VI, preferentially responding in leaf and ear tissues, showing mainly down-regulated expression in leaf while mainly up-regulated expression in ears and tassels; *Mo-7,* consisting of Clade II, where 80% (24/30) of members show no drought response, with the vast majority absent from collinear networks; *Mo-8,* consisting of Clade VI, preferentially responding in ears and leaf tissues with mostly down-regulated expression; *Mo-9,* consisting of Clade I, responding only in leaf and ear tissues; *Mo-10,* consisting of Clade I, preferentially responding in ear tissues; *Mo-11,* consisting of Clade I, preferentially responding to drought in leaf tissues; *Mo-12,* consisting of Clade IX and VII, showing low stress responsiveness; *Mo-13,* consisting of Clade X and XI, where members from different clades exhibit distinct expression patterns: only one Clade X member responds transiently in ears, while all Clade XI members show high stress responsiveness across multiple tissues and stages.

Members of different ancestral genes exhibit differentiation in their response patterns: this is manifested in stress response intensity, as seen in *Mo-7*, where most members show no response; in distinct tissue-specific response preferences, such as *Mo-6* and *Mo-9*, which exhibit high response preference in leave and ear but low or no response in tassels; and in stress response regulatory patterns, such as *Mo-8* and *Mo-6,* where the vast majority of members show down-regulated expression in leaf tissue.

### The analysis of the transcriptional activity s and subcellular localization of *NAC* gene in maize shows functional diversification

Most of the *NAC* across all the plant species analyzed were predicted to be nuclear localized. In order to examine the subcellular localization of *ZmNAC* its full-length coding the *35S:ZmNAC-GFP* fusion proteins were transiently expressed in maize leaf protoplasts. As a control, the fluorescence signal from GFP was detected in both the cytosol and the nuclei, whereas the fluorescence of the *35S:ZmNAC-GFP* fusion proteins was only detected in the nucleus which are in agreement with their role of TFs. Subcellular localization analysis revealed that all 14 *ZmNACs* were localized to the nucleus (Fig. 6a). This aligns with prior reports, demonstrating the evolutionary conservation of *NAC* transcription factor functions. *NAC* transcription factors bind to *NACRS* elements to activate or repress the expression of downstream target genes. When regulating promoters containing *NACRS* elements, *NACs* function as transcriptional activators or repressors. To investigate this dual regulatory capability, we performed dual luciferase (LUC) reporter assays. Triple-tandem *NACRS* sequences were synthesized and inserted upstream of the *CaMV35S* minimal promoter, drving the firefly LUC gene. The Renilla luciferase (rLUC) gene driven by the *CaMV35S* promoter was co-expressed as an internal control. The full-length *ZmNACs*, whose expression was driven by the *CaMV 35S* promoter, were used as effectors. *GFP* served as the control (Fig. 6b). The results showed that Zm00001d002285, Zm00001d003626, Zm00001d016950, Zm00001d018038, Zm00001d036364, Zm00001d047554 and Zm00001d050960 significantly activated the expression of LUC. Conversely, Zm00001d008399, Zm00001d022424, Zm00001d028995, Zm00001d028999 and Zm00001d049860 significantly inhibited the expression of LUC. Notably, Zm00001d027530 and Zm00001d006106 may not specifically recognize the cis-acting elements of *NACRS* (Fig. 6c).

**Figure. 6.**
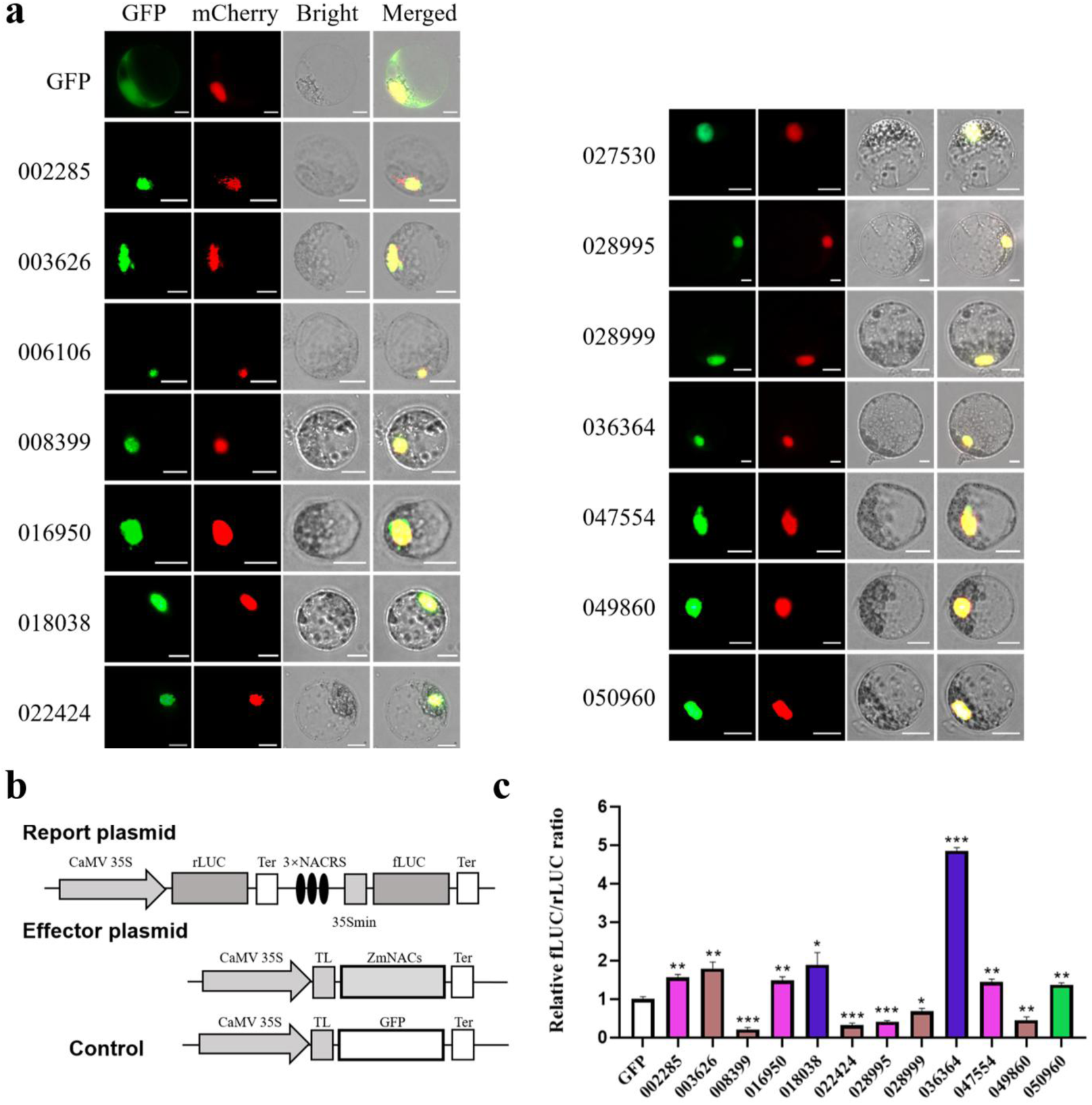
Subcellular localization of ZmNAC proteins, Assay of transcriptional regulation of ZmNACs through dual LUC reporter assay. (a) The fusion protein *P163-ZmNAC-GFP* and *P163-GFP* (control) constructs were transiently expressed in maize B73 protoplasts and visualized with fluorescence microscope after 16h. Nuclei are shown with DAPI staining. Bars=10 μm. (b) CaMV 35S promoter driving *ZmNACs* were used as the effector, and the GFP expression vector was used as a control. The dual luciferase reporter construct consists of 35S driving the Renilla luciferase (rLUC) reporter gene for internal normalization, and triple-tandem *NACRS* sequence driving the firefly luciferase (fLUC) reporter gene. Ter, terminatior sequence; TL, translational leader sequence. (c) ZmNACs showed different activities in dual LUC assay. Data are the mean (±SD)of at least three biological replicates. Asterisks denote significant differences by Student’s t-test (two-tailed, *P<0.05, **P<0.01, ***P<0.001).

Genes derived from the same ancestral gene exhibit conserved transcriptional regulatory activities. Specifically, Zm00001d008399, Zm00001d022424, and Zm00001d028999, which originate from ancestral *Mo-11,* all demonstrate transcriptional repressive activity through *NACRS* element binding, whereas Zm00001d016950 and Zm00001d047554 from ancestral *Mo-1* display transcriptional activation properties. Notably, Zm00001d028995 a paralog sharing both the ancestral origin *Mo-1* and community 42 with Zm00001d047554 exhibits transcriptional repressive activity. This divergence underscores functional differentiation within homologous gene clusters, revealing hierarchical subfunctionalization among ancestral lineages.

### Overexpression of maize *NAC* genes in Arabidopsis regulates the stress resistance of plants

*ZmNACs* widely respond to stress **(Fig.7d)**, which indicates that *ZmNACs* might be involved in stress tolerance. To elucidate the specific function of *ZmNACs*, eight transgenic Arabidopsis plants overexpressing *ZmNACs* gene were generated.

**Figure. 7.**
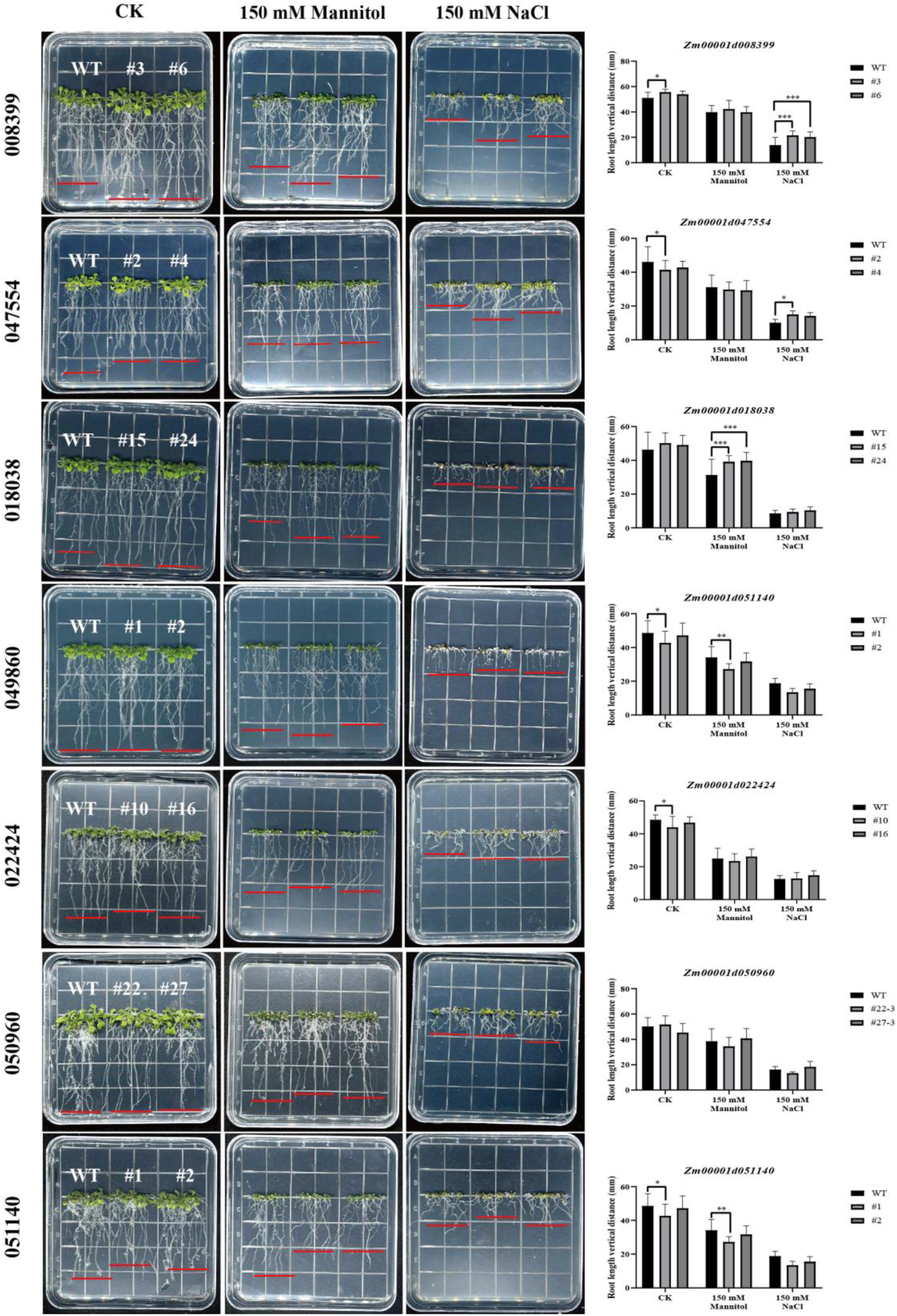
The phenotypes of maize *NAC* overexpressing Arabidopsis under drought and salt stress. (a) Root growth phenotypes and quantitative comparison of root length (b) Col-0 and maize *NAC* overexpressing Arabidopsis on 1/2 MS medium containing 0, 150 mM mannitol, and 150 mM NaCl, respectively. Data represent the mean values (±SD) of at least three biological replicates. Asterisks indicate significant differences by Student’s t-test (*P<0.05, **P<0.01, ***P<0.001).

Assays on the drought and salt tolerance of *ZmNACs* overexpressing in Arabidopsis were conducted using transgenic lines. To determine the root length of *ZmNACs* transgenic lines under drought and salt stress, 1/2MS medium supplemented with 150 mM mannitol and 150 mM NaCl was used to simulate drought and salt stress conditions. Homozygous transgenic Arabidopsis plants were selected, germinated for three days, and then transferred to the simulated drought and salt stress environments; nine days later, the root lengths of the transgenic lines and wild-type plants were calculated. Under normal condition, *Zm00001d008399 (Mo-11)* overexpressing Arabidopsis exhibited longer roots than wild-type plants. Under drought stress, there was no significant difference between the overexpression lines and wild-type plants. Under salt stress condition, the roots of overexpressing lines were significantly longer than those of wild-type plants. Thus, *Zm00001d008399* shows insensitivity to drought but significantly enhances salt tolerance. Under normal condition, *Zm00001d047554 (Mo-1)* overexpressing Arabidopsis lines had slightly shorter roots than wild-type plants. Under drought stress, there was no difference in root length between the overexpression lines and wild-type plants. Under salt stress, the roots of *Zm00001d047554* overexpressing lines were significantly longer than those of wild-type plants. Therefore, *Zm00001d047554* improves drought resistance and significantly enhances salt tolerance in transgenic overexpression Arabidopsis. Under normal and salt stress conditions, the roots of *Zm00001d018038 (Mo-6)* overexpressing lines showed no difference from thoese of wild-type plants. Under drought stress, the roots of *Zm00001d018038* overexpressing lines were significantly longer than those of wild-type plants. This indicates that *Zm00001d018038* enhances drought resistance in transgenic overexpression Arabidopsis while being insensitive to salt stress. *Zm00001d049860 (Mo-9)* overexpressing Arabidopsis plants showed no significant differences from wild-type plants under normal, drought, and salt stress conditions, indicating that *Zm00001d049860* is insensitive to both drought and salt stress. Under normal conditions, the roots of *Zm00001d022424 (Mo-11)* overexpressing lines were shorter than those of wild-type plants. Under drought stress, there was no difference in root length between the overexpression lines and wild-type plants. Under salt stress, the roots of *Zm00001d022424* overexpressing lines were significantly longer than those of wild-type plants. Consequently, *Zm00001d022424* enhances drought resistance and improves salt tolerance in transgenic overexpression Arabidopsis. Under normal, drought, and salt stress conditions, the roots of *Zm00001d050960 (Mo-4)* overexpressing lines showed no difference compared to those of wild-type plants, suggesting that *Zm00001d050960* is insensitive to both drought and salt stress. Under normal and drought stress conditions, the roots of *Zm00001d051140 (Mo-4)* overexpressing lines were significantly shorter than those of wild-type plants; no difference under salt stress conditions. Thus, *Zm00001d051140* is insensitive to drought stress but improves salt resistance.

These results indicate that maize *NAC* genes within the same clade exhibit distinct response patterns to drought and salt stress, and functional divergence in stress resistance has emerged among *NAC* genes derived from different ancestral lineages.

## Discussion

### Rapid genome sequencing accelerates the comprehensive evolutionary investigation of NAC proteins in plants

Numerous studies on the phylogenetic analysis of gene families have been reported in recent years; however, most have focused solely on single species, providing only limited evolutionary insights. With the completion of genome sequencing for more species, an increasing number of researchers have expanded their studies to multiple species (Artur et al., 2019; Lama et al., 2019; Tan et al., 2018). In the present study, to shed light on the evolutionary mechanisms and reveal the dynamic changes of *NAC* genes in the evolutionary process, we performed genome-wide analyses of *NAC* genes from 4 major land plant lineages, and the NAC proteins of 17 species were not identified previously. In total, 5,052 *NACs* were identified across 46 species, and the differential counts partially reflected the complex patterns of gene loss or WGDs. For example, the number of *NAC* genes in *Glycine max* is 176, approximately twice taht in *Phaseolus vulgaris* and *Cajanus cajan*, and this may be caused by a WGD of soybean genome (Schmutz et al., 2010). Generally, in angiosperms, the number of *NACs* in monocots is relatively conservative (ranging from 75 in Orchid to 165 in Banana), while that in dicots has a larger range (ranging from 48 in Sugar beet to 226 in Apple) (Fig. 1).

The phylogenetic analysis 5,052 *NACs* grouped 11 evolutionary clades into three categories, being consistent with specific conserved motifs of NAC proteins in different clades. This suggests that the evolution of *NAC* genes coincides with the development of significant plant lineage. Furthermore, numerous previous studies have reported that genes within the same clade often perform similar functions. For example, Arabidopsis *CUC1/2, NAC1, At5g07680*, *At5g61430,* and Rice OMTN1-6, which are clustered in Clade Ⅶ, are all regulated by *miR164* (Furuta et al., 2014; Guo et al., 2005; H. Kim et al., 2009; Laufs et al., 2004). This indicates that the expression regulation patterns of genes from different species are conserved within the same clade. Similarly, most secondary wall NACs (SWNs), which act as master regulators of secondary cell wall development in vascular tissues(Handakumbura & Hazen, 2012; Zhong et al., 2006), are localized in two land plant-specific clades (Clades Ⅶ and Ⅷ). These NAC subdomains are highly conserved (Fig. S1), suggesting that the genes within these two clades may maintain the potential function associated with secondary cell wall development.

### Synteny network approach advances the understanding of the evolutionary relationships between NAC proteins in plants

In this study, we totally identified 23.3% (1,177/5,052) tandem duplications of *NAC* genes. This suggests that tandem duplication events are one of the factors driving *NAC* family expansion. We found that there are a wide range of tandem duplications between certain clades, and the partners of the tandems displayed similar and conserved differential sites. The synteny network analysis confirmed that there are strong synteny relationships between these tandems, meaning these tandems have occurred ancient tandem gene arrangement in angiosperms.

Furthermore, based on the synteny network, we also identified some specific evolutionary events within communities, such as lineage-specific transpositions. In *NAC* genes, especially in four angiosperm-specific clades, all communities are lineage-specific, suggesting the expansion of *NAC* genes may be accompanied with the divergence of evolutionary lineages and ancient polyploidy events. Other 34 communities spread widely in angiosperms and include nearly half of the *NAC* genes (44.6%; 2,255/5,052), so we reckoned that the syntelogs of these angiosperm-wide communities retained more conserved evolutionary relationships and also were accompanied with WGD or segmental duplication events in evolutionary process.

### Multi-omics data integrative analysis reveals functional diversification of *NAC* gene family

The maize *NAC* genes trace back to 13 ancestral *NAC* genes originating at distinct evolutionary nodes in plants. Tissue-specific expression and stress-response pattern analyses demonstrated significant expression divergence of *ZmNACs* both between and within clades, revealing differential expansion degrees and functional fates of ancestral genes during evolution(Fig. 5). NAC transcription factors are widely reported to localize to the nucleus, and all 14 tested *ZmNACs* in this study showed nuclear subcellular localization, confirming the conserved nuclear functionality of *NACs* as transcription factors. Transcriptional activity assays revealed relative consistency within clades but significant divergence between clades, which reflects functional differentiation and diversity in *NAC* transcriptional regulation (Fig. 6). Although we traced the emergence timeline of these genes within land plant lineages, pinpointing the acquisition of specific functions remains challenging. Specific transcription factor functions might have predated duplication events but were subsequently lost in some paralogs, or alternatively, emerged post-duplication through neofunctionalization. *ZmNACs* clustered in collinearity networks exhibited widespread cross-species synteny, suggesting evolutionary retention and conserved functions in land plants. Expression profiling and drought-stress response analyses revealed that most syntenic *ZmNACs* are expressed and responsive to drought stress. Functional validation using transgenic Arabidopsis materials under abiotic stress confirmed their regulatory roles in stress responses, with different genes exhibiting functional diversity in stress modulation.

By identifying over 5,000 *NAC* genes across 46 land plant species, we successfully reconstructed the phylogenetic framework of land plant *NAC* genes. Thirteen ancestral *NAC* genes were identified in land plant ancestors, with *NAC* family expansion predominantly occurring after angiosperm divergence despite varied evolutionary trajectories. Our study facilitates the functional elucidation of *NAC* genes across different plant species and provides new insights into the evolution of plant gene families.

### Conclusions

In the present study, phylogenetic analysis of *NAC* transcription factors across 46 land plants was carried out to explore the contribution of gene replication events to the expansion of *NAC* gene family, and the evolution of tandemly replication and membrane-localized *NAC* genes was performed by using collinearity networks, by reconstructing the evolutionary histories of *NAC* genes in maize and Arabidopsis, we investigated their distinct evolutionary fates. Subsequently, combined with the RNA-seq analysis of different growth and development stages and drought stress treatments, the subcellular localization, transcriptional activity and function of potential candidate *NAC* genes in maize were studied. A variety of approaches was applied to identify the evolutionary processes likely contributing to divergence within the gene family. The findings indicate multiple levels of functional divergence(Fig. 8). These results contribute to the functional characterization of the *NAC* gene family, and the genes identified in this study related to salt and drought stress responses may serve as potential genes for maize improvement.

**Figure. 8.**
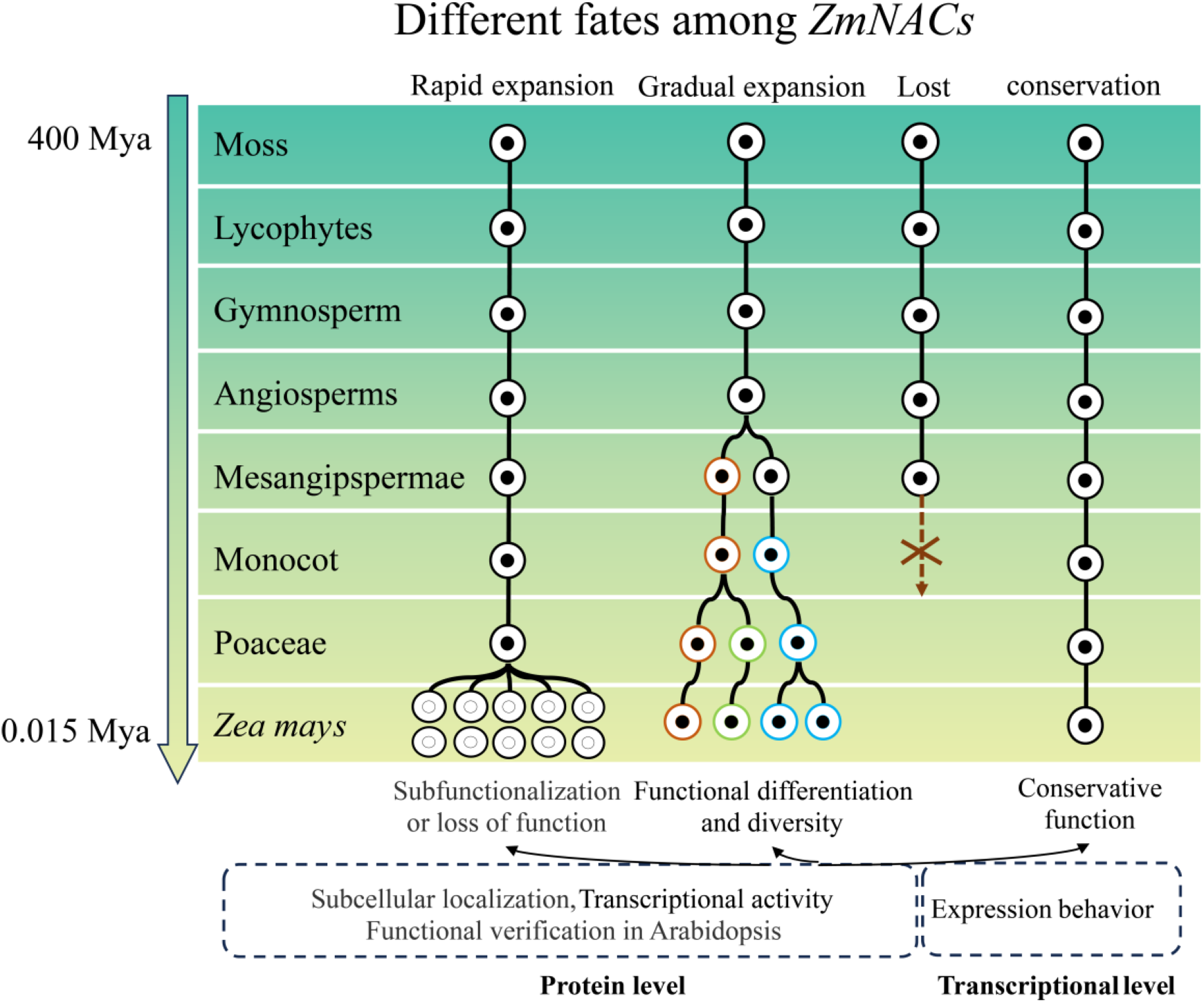
Different evolution fates among *ZmNACs*

## Materials and methods

### Identification of NAC proteins in 46 land plant genomes

We performed the phylogenetic analysis of NAC proteins across 46 land plants, encompassing moss, lycophytes, gymnosperms, early angiosperms, monocots, early eudicots, asterids, and rosids (Fig. 1; Supporting Information Table S1). The genomic sequences, annotations, and gene models for maize *(Zea mays*) and cucumber *(Cucumis sativus*) were procured from the Ensembl Plants database (http://plants.ensembl.org/index.html) (Kersey et al., 2018), and those for the other 44 species were obtained from Zhao et al. (Zhao et al., 2017).

For each of these 46 genomes, putative NAC proteins were initially identified by scanning against the corresponding protein sequence using HMMER v3.0 software (Eddy, 2011) with an threshold of E-value < 0.001, using Hidden Markov Model (HMM) profile of the *NAC* domain (Pfam: PF02365) (http://pfam.xfam.org) (El-Gebali et al., 2019). Secondly, the putative NAC proteins were further filtered by removing those with an *NAC* domain sequence shorter than 80 residues.

### Sequence alignment and phylogenetic analysis

Multiple sequence alignment of all predicted NAC proteins was performed using the DNAMAN v5.0 and MAFFT program (http://mafft.cbrc.jp/alignment/software) (Katoh & Standley, 2013). The phylogenetic tree was constructed based on the resulting alignments using the FastTree software (http://www.microbesonline.org/fasttree) (Price et al., 2010). The stability of internal nodes was assessed by bootstrap analysis with 1,000 replicates. A zygnematophycean NAC protein from *Mesotaenium endlicherianum* was used as the outgroup. The phylogenetic tree was annotated and visualized using iTOL (Letunic & Bork, 2016) and FigTree v1.4.3 program (http://tree.bio.ed.ac.uk/software/figtree/).

Conserved motifs within the *NAC* domain were identified using the MEME v5.1 program (http://meme.sdsc.edu) (Bailey et al., 2009), and the generated sequence logos were manually curated to compare their differences among different clades. The membrane-bound NAC proteins (NTLs) were predicted using the TMHHM server v2.0 (http://www.cbs.dtu.dk/services/TMHMM/).

Gene gains and losses at each divergence node were reconciled using Notung software (Stolzer et al., 2012). The number of ancestral *NAC* genes was determined using monophyletic *NAC* lineages as described in Shao et al.(Shao et al., 2014). The exon–intron structures were visualized using the Gene Structure Display Server (GSDS, http://gsds.cbi.pku.edu.cn) (Hu et al., 2015).

### Tandem duplication and synteny network analysis of *NAC* gene family

Tandem duplications were characterized by generating nearby gene copies as multiple members of this family (Cannon et al., 2004). The tandem duplication of paralogous *NAC* genes was defined as those closely related genes falling within 100 kb of one another and separated by ten or fewer genes.

We used the Synets method (Zhao et al., 2017; Zhao & Schranz, 2017) for syntenic block calculation and network construction (https://github.com/zhaotao1987/SynNet-Pipeline). Briefly, pairwise whole-genome comparisons were performed using DIAMOND (Buchfink et al., 2014). Synteny block detection was performed with MCScanX software with its default parameters (Wang et al., 2012). The syntenic blocks containing the identified *NAC* sequences were used to build synteny networks that were visualized with Cytoscape 3.6.0 (Shannon et al., 2003) and Gephi 0.9.1(Bastian et al., 2009). The HC-PIN clustering algorithm in CytoCluster (Li et al., 2017) was selected to perform the synteny network clustering, employing the strong setting and a complex size threshold of 4. Subsequently, synteny communities derived from CytoCluster were extracted and converted into a heatmap by using the ‘pheatmap’ package in the R programming language (https://www.r-project.org).

### Reconstruction of Ancestral *NAC* Genes

The phylogenetic analysis was conducted by executing multiple sequence alignment (MAFFT) of *NAC* sequences derived from Arabidopsis thaliana and A. lyrata (genus Arabidopsis). These sequences were aligned with those obtained from the outgroup species, Brassica oleracea, as well as the NAM domain of the outgroup gene *Carica papaya NACs*. The alignment results served as the foundation for constructing a phylogenetic tree. Subsequent analysis of the resulting gene and species trees was performed using Notung software to infer events of gene duplication and loss. Additionally, the origin time of each *NAC* branch was estimated based on the structure of the Notung-reconciled tree. In this context, if a branch of *NAC* genes on the tree is monophyletic and its origin time can be traced back to the common ancestor of the genus Arabidopsis, then this branch is classified as an *NAC* gene family of that genus. Conversely, if the origin time of *NAC* genes in a monophyletic group is traced back to the common ancestor of both Arabidopsis and Brassica (*family Brassicaceae*), then this branch is defined as a Brassicaceae *NAC* gene family. This systematic approach was applied to infer the evolutionary history down to *Physcomitrium patens*.

### Tissue-specific and stress-related expression patterns of *NAC* genes in maize

The RNA-Seq data from B73, encompassing six tissues including leaf, shoot, seedling, bract, pericarp, and silk were used to analyze tissue-specific expression profile of *ZmNACs* (Wang et al., 2018). The expression profiles of *NAC* genes in maize were examined using stress-related RNA-seq data from three tissues (leaf, ear, and tassel) across four developmental stages (V12, V14, V18, and R1) under two environmental conditions (well-watered and drought)(Thatcher et al., 2016). Low-quality reads and sequencing adapters were trimmed using fastp with default parameters (Chen et al., 2018). Subsequently, clean reads were aligned to the B73 reference genome using HISAT2 (Bastian et al., 2009) with the parameters (min-intronlen=20, max-intronlen=10000, k=20) (Chen et al., 2019). The transcript expression level of *NAC* genes was evaluated using uniquely-mapped reads extracted from each alignment result as input to StringTie (Pertea et al., 2015) with reference genome annotation (APGv4.33) in term of FPKM (Fragments Per Kilobase of transcript per Million fragments mapped). The RNA-seq and ribosome profiling data were retrieved from the SRA database (Accession number: SRP052520) (Lei et al., 2015) and processed following the procedure described in Tan *et al*.(Tan et al., 2018).

### Plant materials and Growth Conditions

The genetic background used in our study was *Arabidopsis thaliana* ecotype *Columbia-0* (Col-0). The overexpression vectors were constructed by inserting the coding sequences of *ZmNACs* between the *CaMV35S* promoter and the nos terminator in *pBI-III*. The *Agrobacterium tumefaciens*-mediated floral-dip method was used to obtain transgenic Arabidopsis plants. Seeds were sterilized and stratified at 4°C for 3 days in darkness, and then germinated on 1/2MS medium at 22℃, 60–80% relative humidity under a 16 h light/8 h dark cycle. The maize inbred line B73 seeds were imbibed overnight on sterilized filter paper soaked with sterile water, then planted in vermiculite-filled pots and grown at 28°C for two weeks under a 16 h light/8 h dark cycle. The mesophyll tissue of B73 seedlings was utilized to isolate Maize protoplasts. *Nicotiana benthamiana* plants were cultivated at 25°C for 5-6 weeks under a 12 h light/12 h dark cycle and their leaves were used for infiltration.

### Subcellular localization of ZmNAC proteins

The coding regions of *ZmNACs* were amplified using PCR with gene-specific primers and independently fused to the N-terminus of GFP in the *p163* vector. Maize protoplasts were isolated from the mesophyll tissue of 2-week-old maize seedings grown under a 16h light/8h dark cycle and then transformed using the PEG transfection method (Yoo et al., 2007). After incubated in darkness at 25℃ for 16 h, the GFP fluorescent signals were monitored using a laser-scanning confocal microscope (DM5000 B, Leica, Germany). The *NLS-mCherry* marker indicates the nuclei (Yan et al., 2021).

### Dual luciferase reporter assay

The CDSs of *ZmNACs* were cloned into the *pBamEYFP* plasmid and served as effectors. The reporter plasmid *3×NACRS::LUC* contains the tandem repeats of the triple-tandem NACRS element and a 46-bp *CaMV35S* minimal promoter (Chen et al., 2023). The control plasmid was the binary vector pBamEYFP, in which the *EYFP* gene was driven by the *35S* promoter of *CaMV*. The agrobacterial cultures transformed with the effector and reporter plasmid (9:1, v/v) were co-infiltrated into N. benthamiana leaves (Chen et al., 2023). Samples were harvested at 72 h post-transfection. The LUC and REN activities were determined using a dual-luciferase reporter gene assay kit (Yeasen, China), with three biological replicates were assayed.

### Drought and salt tolerance assays of transgenic plants

Three-days-old seedlings were transferred from 1/2MS to 1/2MS medium supplemented with 0, 150 mM Mannitol, or 150mM NaCl and grown vertically. The plants were photographed after 9 days, and the root lengths were measured using ImageJ software. Three independent experiments were performed and the data from each experiment was used for statistical analysis. The Student’s t-test was employed to evaluate the differences between the Col-0 and transgenic plants.

## Data availability

All data generated or analyzed during this study are included in this published article and its supplementary information files.

## Abbreviations

*RNA*: Ribonucleic acid
*DNA*: Deoxyribonucleic acid
*WT*: Wide Type
*DEGs*: Differentially expressed genes
*N. benthamiana*: *Nicotiana benthamiana*
*GFP*: Green Fluorescent Protein
*EYFP*: Enhanced yellow fluorescent protein (YFP)
*PCR*: Polymerase Chain Reaction
*ANOVA*: One-way analysis of variance

## Conflict of interest

The authors declare that they have no conflict of interest.

## Author contributions

YL, ZL and YZ conceived and designed the research; YL and YW performed the research; YL, YZ, LG and XC performed the data analysis; ZL, YZ and LG performed the interpretation of results; YL, ZL and LG wrote the manuscript. All authors proofread the final version of the manuscript.

## Acknowledgements

This work was supported by the National Natural Science Foundation of China (31201268), the Special Fund for Basic Scientific Research of Central College (2452019072).

## Supplemental Data

**Fig. S1 Phylogenetic tree of 5052 NAC proteins from 46 species**. The phylogeny was constructed using the conserved NAC domain. The NAC proteins were divided into 11 clades (Ⅰ, Ⅱ, Ⅲ, …Ⅺ) shown with different colors except 8 NACs which conflicted with the overall tree topology (black branches). The percentage of species categories for each clade was represented as pie chart and colored according to the species phylogeny (Table S3 and Fig. 1). The black circle marked the four NTLs from *P. patens*.

**Fig. S2 The conserved motifs of NAC domain among 11 clades**. Subdomains A to E are shown by dotted line above the sequences. The bit score indicates the information content for each position in the sequence.

**Fig. S3 The pie chart presents the percentage of NTMs in each clade**. (**a)** The presents of NTMs containing one α-helical membrane-bound domain. (**b)** The presents of NTMs containing more than one α-helical membrane-bound domains.

**Fig. S4 Summary of the occurrence of seven inferred ancient tandem gene arrangements detected in the species analyzed**. The species tree on top of the phylogenetic profiling is a simplified version (without species names) of the tree used in Fig 1.

**Fig. S5 Proposed evolutionary scenario for the origin of the *NAC* gene family in community 2.** The orange box and blue box indicate the *NAC* genes from Clade Ⅸ and Clade V, respectively. Black helix indicates membrane-bound domain and black line indicates chromosome.

**Fig. S6 Phylogenetic tree of moss *NAC* genes**. The lines beside the phylogenetic tree indicate the monophyletic NAC lineages. The black circle indicates the NTLs.

**Fig. S7 Phylogenetic tree of lycophyte *NAC* genes.**

**Fig. S8 Phylogenetic tree of gymnosperm *NAC* genes.**

**Fig. S9 Phylogenetic tree of angiosperm *NAC* genes.**

**Fig. S10 Phylogenetic tree of monocot *NAC* genes.**

**Fig. S11 Phylogenetic tree of dicot *NAC* genes.**

**Fig. S12 Phylogenetic tree of rosid *NAC* genes.**

**Fig. S13 Phylogenetic tree of brassicaceae *NAC* genes.**

**Table S1. The putative *NAC* genes in 46 species.**

**Table S2. Number of NAC proteins found in 46 species.**

**Table S3. The percentage of species categories for each clade.**

**Table S4. Putative membrane-bound *NAC* genes in 46 species.**

**Table S5. The tandem duplications found in 46 species.**

**Table S6. The syntenic gene pairs of *NAC* gene family.**

**Table S7. The 95 communities of syntenic *NAC* genes.**

**Table S8. Ancestral *NAC* genes expand to the current Arabidopsis and Maize *NAC* genes.**

**Table S9. The expression of *ZmNAC* in six tissues.**

**Table S10. Ka/Ks value of tandem duplications in Clade Ⅱ.**

**Table S11. Differential expression level (Log2Foldchange) of maize NAC genes in three tissues.** (Ear, Leaf, Tassel) at four developmental stages (V12, V14, V18) of maize under well-watered (WW) and drought stress (DS) conditions.

## Notes

### Competing Interest Statement

The authors have declared no competing interest.

