## Supplemental Figure S1-S13 for "Insights into the patterns of molecular evolution and functional diversification of *NAC* gene family in land plants"

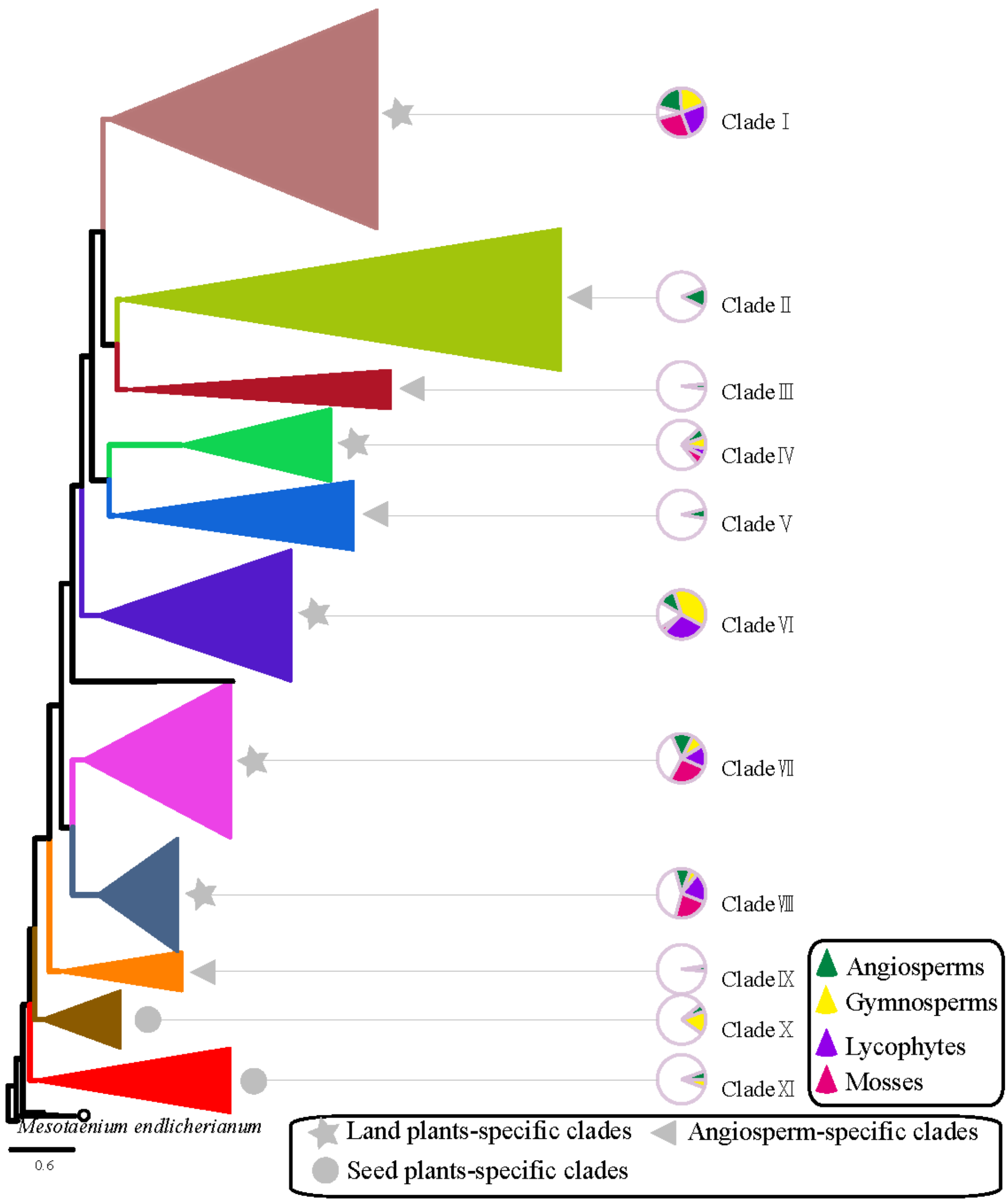


**Fig. S1 Phylogenetic tree of 5052 NAC proteins from 46 species**. The phylogeny was constructed using the conserved NAC domain. The NAC proteins were divided into 11 clades (I, II, III, …XI) shown with different colors except 8 NACs which conflicted with the overall tree topology (black branches). The percentage of species categories for each clade was represented as pie chart and colored according to the species phylogeny (Table S3 and Fig. 1). The black circle marked the four NTLs from *P. patens*.

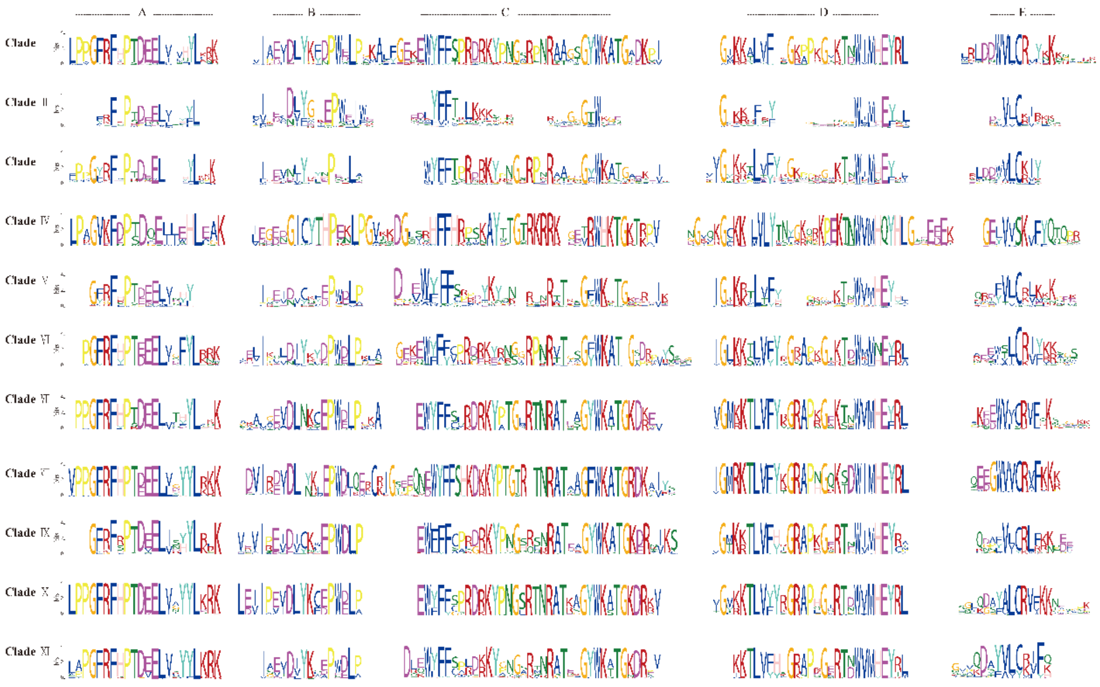

**Fig. S2 The conserved motifs of NAC domain among 11 clades**. Subdomains A to E are shown by dotted line above the sequences. The bit score indicates the information content for each position in the sequence.

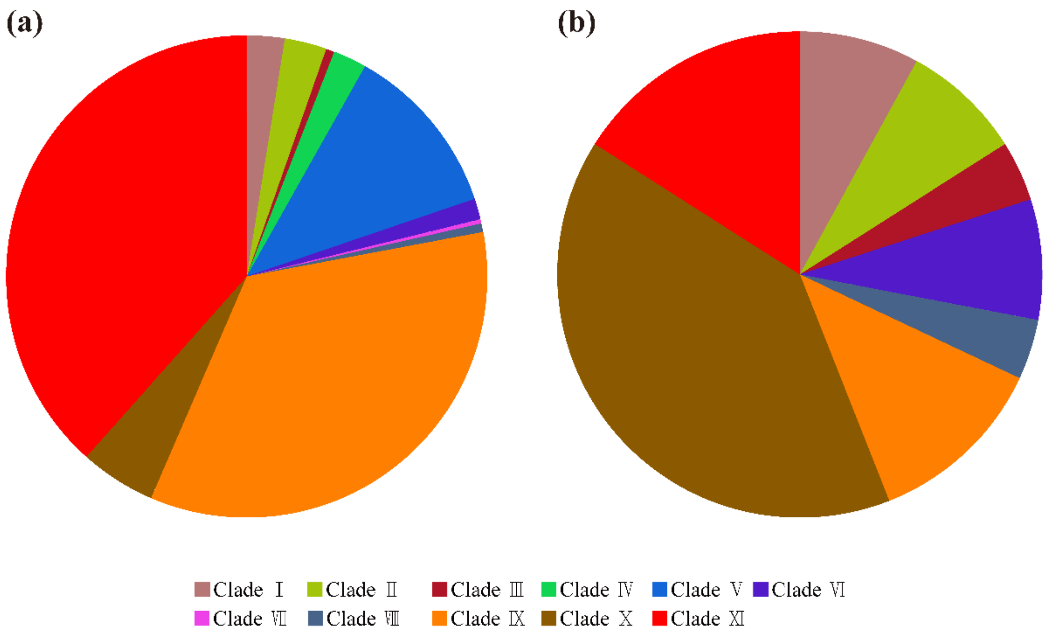


**Fig. S3 The pie chart presents the percentage of NTMs in each clade**. (**a)** The presents of NTMs containing one α-helical membrane-bound domain. (**b)** The presents of NTMs containing more than one α-helical membrane-bound domains.

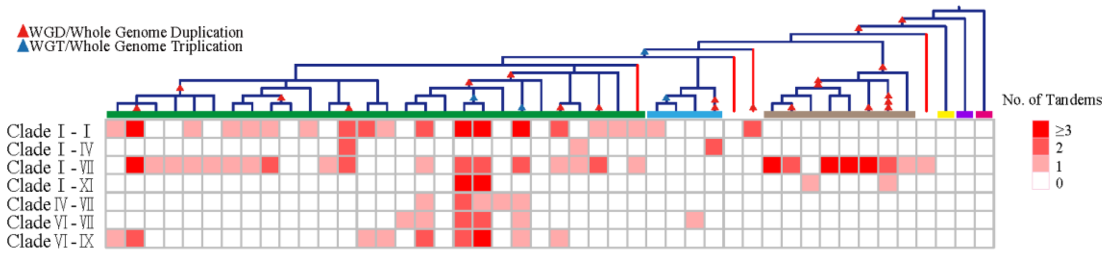


**Fig. S4 Summary of the occurrence of seven inferred ancient tandem gene arrangements detected in the species analyzed**. The species tree on top of the phylogenetic profiling is a simplified version (without species names) of the tree used in Fig 1.

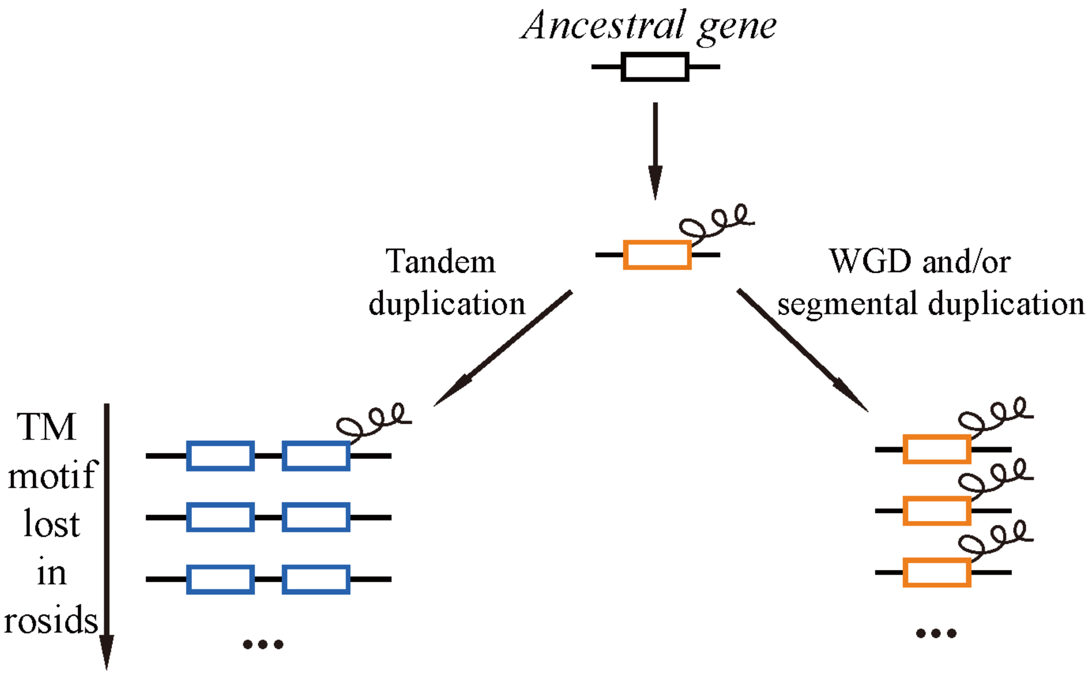


**Fig. S5** Proposed evolutionary scenario for the origin of the *NAC* gene family in community.

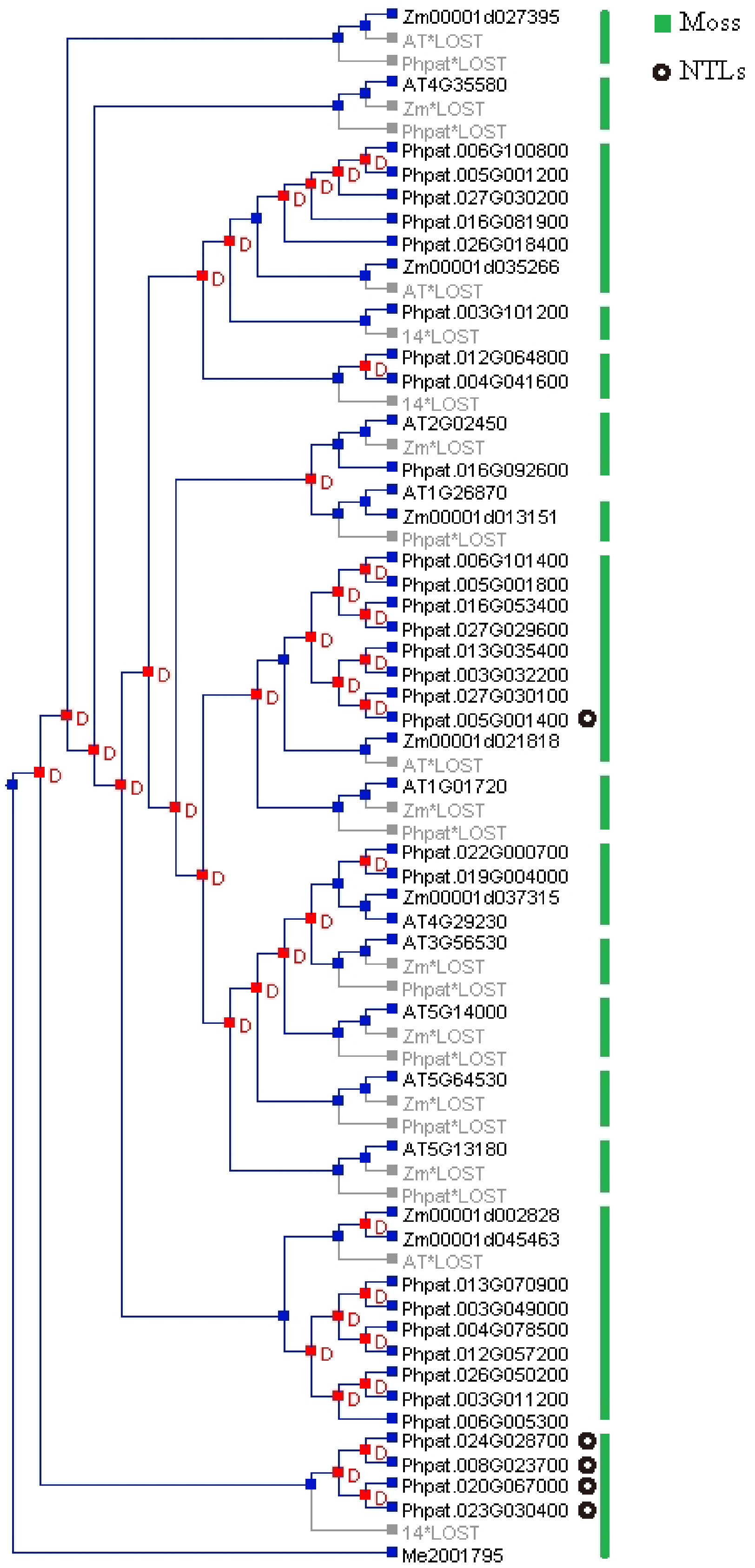


Image generated with Notung 2.9.1.3, on 2019-12-17

**Fig. S6** Phylogenetic tree of moss *NAC* genes.

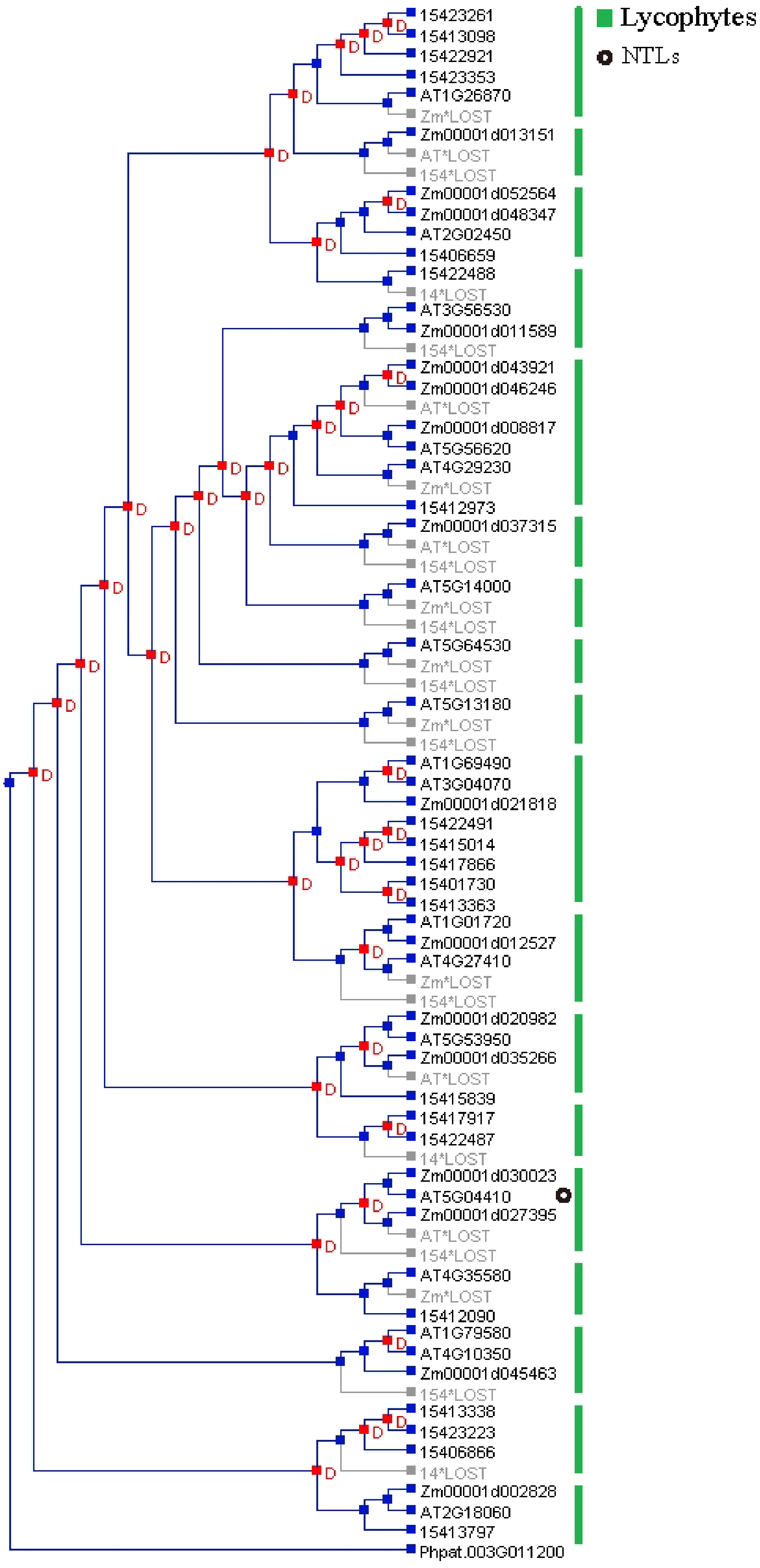


Image generated with Notung 2.9.1.3, on 2019-12-16

**Fig. S7** Phylogenetic tree of lycophyte *NAC* genes.

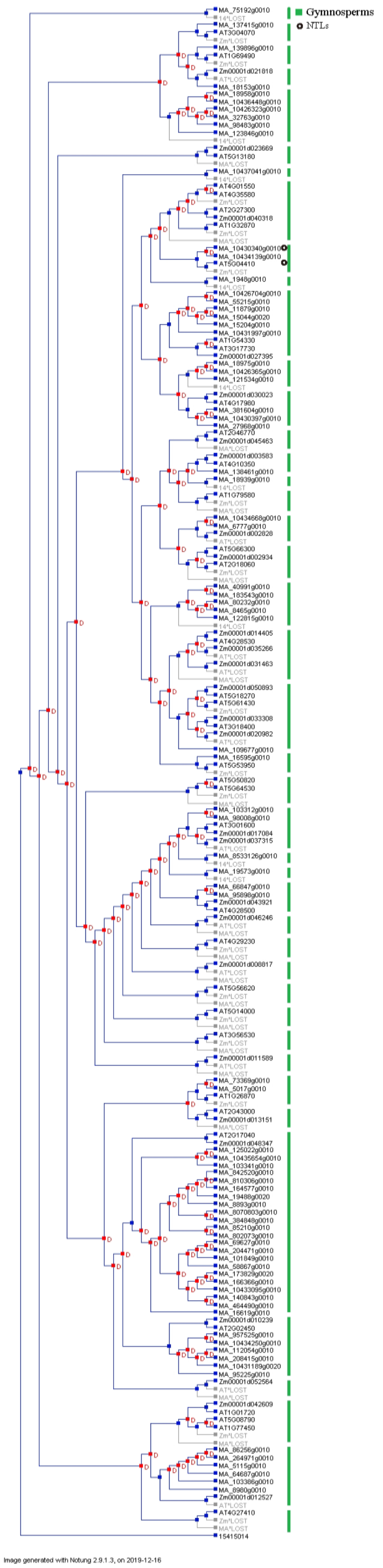


**Fig. S8** Phylogenetic tree of gymnosperm *NAC* genes.

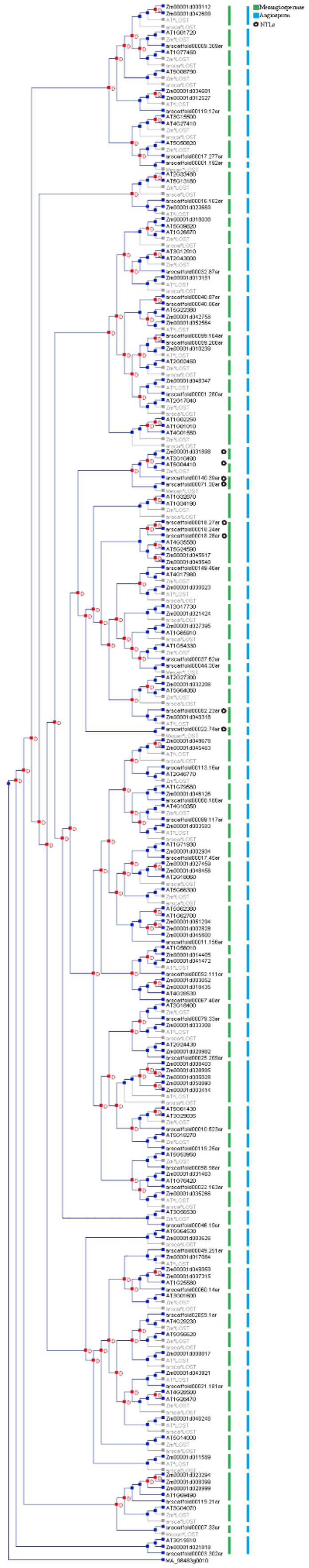

Image generated with Notung 2.9.1.3, on 2019-12-10

**Fig. S9** Phylogenetic tree of angiosperm *NAC* genes.

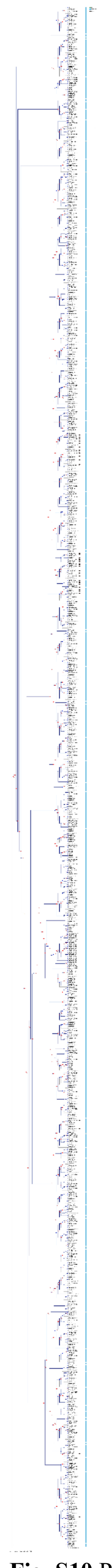

**Fig. S10** Phylogenetic tree of monocot *NAC* genes.

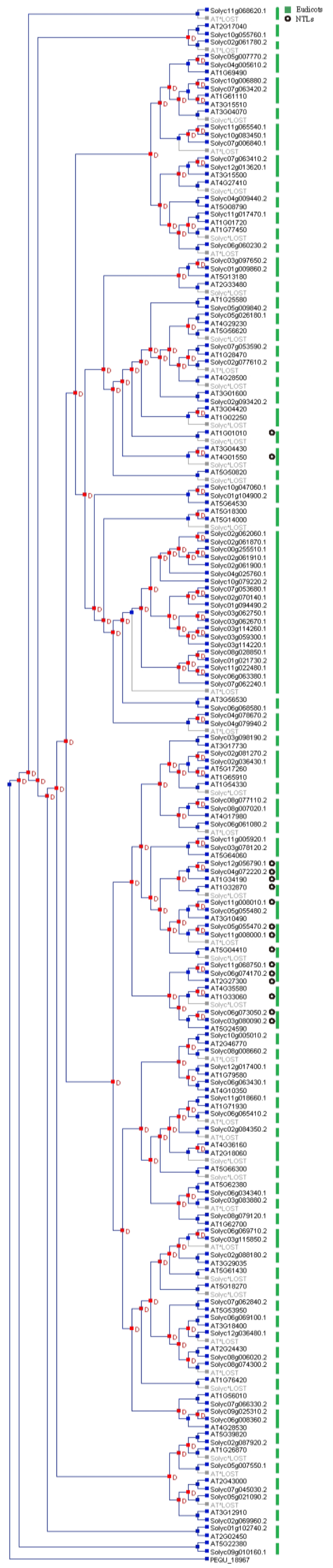


**Fig. S11** Phylogenetic tree of dicot *NAC* genes.

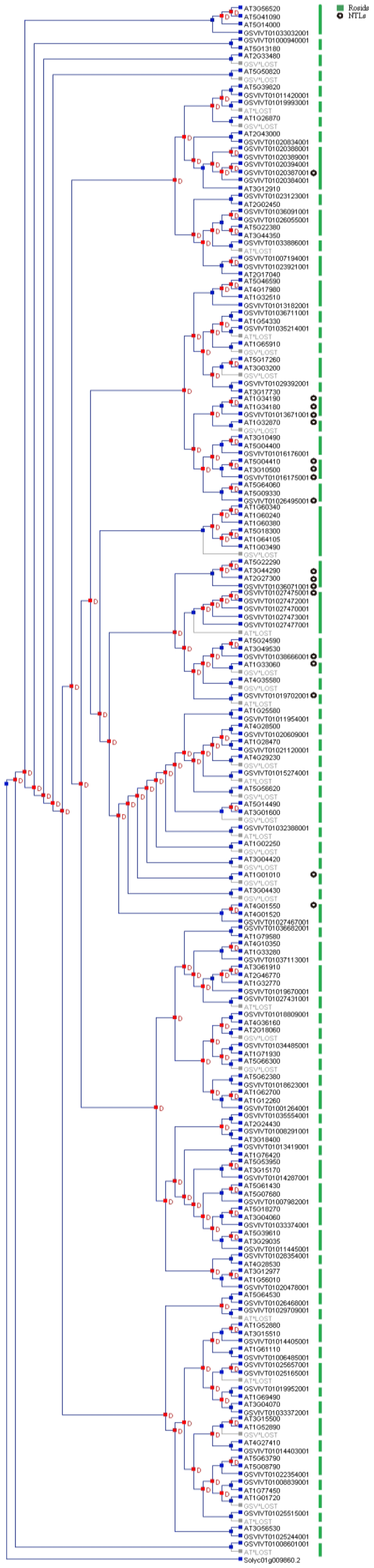


Image generated with Notung 2.9.1.3, on 2019-12-9

**Fig. S12** Phylogenetic tree of rosid *NAC* genes.

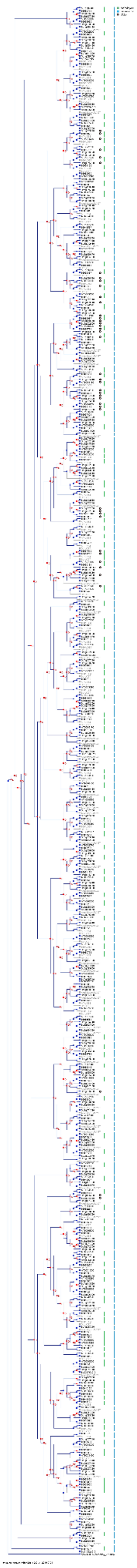

**Fig. S13** Phylogenetic tree of brassicaceae *NAC* genes.
